# A non-canonical lysosome biogenesis pathway generates Golgi-associated lysosomes during epidermal differentiation

**DOI:** 10.1101/312033

**Authors:** Sarmistha Mahanty, Shruthi Shirur Dakappa, Rezwan Shariff, Saloni Patel, Mruthyunjaya Mathapathi Swamy, Amitabha Majumdar, Subba Rao Gangi Setty

## Abstract

Keratinocytes maintain epidermis integrity and function including physical and antimicrobial barrier through cellular differentiation. This process is predicted to be controlled by calcium ion gradient and nutritional stress. Keratinocytes undergo proteome changes during differentiation, which enhances the intracellular organelle digestion to sustain the stress conditions. However, the molecular mechanism between epidermal differentiation and organelle homeostasis is poorly understood. Here, we used primary neonatal human epidermal keratinocytes to study the link between cellular differentiation, signaling pathways and organelle turnover. Upon addition of calcium chloride (2 mM) to the culture medium, keratinocytes increased their cell size and the expression of differentiation markers. Moreover, differentiated keratinocytes showed enhanced lysosome biogenesis that was dependent on ATF6-arm of UPR signaling but independent of mTOR-MiT/TFE transcription factors. Furthermore, chemical inhibition of mTOR has increased keratinocyte differentiation and relocalized the MiT/TFE TFs to the lysosome membranes, indicating that autophagy activation promotes the epidermal differentiation. Interestingly, differentiation of keratinocytes resulted in dispersal of fragmented Golgi and lysosomes, and the later organelles showed colocalization with Golgi-tethering proteins, suggesting that these lysosomes possibly originated from Golgi, hence named as Golgi-associated lysosomes (GALs). Consistent to this prediction, inhibition of Golgi function using brefeldin A completely abolished the formation of GALs and the keratinocyte differentiation. Thus, ER stress regulates the biogenesis of GALs, which maintains keratinocyte differentiation and epidermal homeostasis.

## Introduction

Human epidermis majorly composed of proliferative and differentiated keratinocytes organized into four distinct layers, that are characterized by specific gene expression and the extent of differentiation^1–3^. These sublayers are predicted to be formed and maintained due to pre-existing calcium gradient between the layers^4–7^. Proliferative keratinocytes of stratum basale gradually undergo differentiation towards the upper layers and terminally differentiated in stratum corneum (cornified envelope) that sheds off and replaced by a new layer and thus maintains epidermal homeostasis. Additionally, during differentiation keratinocytes acquire special functions that are linked to skin’s physical and antimicrobial barrier properties. Moreover, studies have shown that differentiated epidermal layers gradually lose their intracellular organelles including nucleus by increasing macroautophagy^8,9^. However, the molecular mechanism of ionic calcium influence on cellular differentiation and its link to organelle homeostasis and autophagy is poorly understood.

Previous studies have shown that extracellular high Ca^2+^ ion concentration activates PI3K (phosphatidylinositide 3-kinase) pathway following the elevation of intracellular Calcium through IP3R (inositol triphosphate receptor), which potentially induce keratinocyte differentiation^5,6,10–13^. However, the downstream signalling of intracellular calcium in regulating the cell differentiation process is largely unknown. It has been shown that PI3K in connection with other regulatory molecules such as mTORC1 (mammalian target of rapamycin complex 1), AMPK (AMP-activated protein kinase) and AKT (also called PKB, protein kinase B) regulates variety of cellular pathways in response to multiple cellular stresses including high calcium, nutrient/amino acid/energy depletion, oxidative stress etc.^14–20^. These kinases activate transcriptional response to compensate or nullify the stress conditions and some times cells undergo differentiation^20–24^. Similarly, studies have shown that mTORC1-dependent MiT/TFE transcription factor (TF) TFEB regulates lysosome biogenesis and cellular autophagy in many cell types^25–30^ and the differentiation of osteoblasts^24^. Moreover, studies have shown that autophagy is required for epidermal differentiation *in vivo* and also during calcium-induced keratinocytes differentiation *in vitro*^8,9,31,32^. Nevertheless, the TFs involved in modulating autophagy process during keratinocyte differentiation remain unknown. Studies have shown that lysosomes play key role in regulating autophagy incuding its turnover/flux^25,26,28^. But, whether the increased autophagy also requires enhanced lysosome biogenesis during keratinocyte differentiation was not yet been addressed. Interestingly, previous studies have shown the accumulation of lysosomal bodies in the upper layers of epidermis^3,33^. Added to the complexity, cytosolic calcium possibly activate endoplasmic reticulum (ER) stress that further enhance the autophagy and cell survival through unfolded protein response (UPR)^34^. Recent studies have shown that ER stress induces lysosome biogenesis and autophagy involving TFEB/TFE3 TFs in an mTORC1-independent manner^35,36^. Although, studies have shown the cross-regulation between autophagy and intracellular Ca^2+^ signaling^18^, but the role of UPR on lysosome biogenesis is not explored.

In mammalian cell, UPR is sensed by the three ER resident sensory kinases, namely IRE1, ATF6 and PERK. During ER stress, dimerized IRE1 and PERK activate downstram TFs XBP-1 and ATF4 respectively. In contrast, ATF6 translocates to the Golgi and then undergo proteolytic cleavage to generate cytosolic TF. These TFs translocate into the nucleus and regulate multiple groups of gene expression such as chaperons, autophagy, ERAD including apotosis^37^. Studies have also shown that UPR TFs crosstalk each other in regulating the gene expression of different pathways^38^. Moreover, intracellular calcium also modulates the UPR pathway to achieve cellular homeostasis^34^. However, the role of ER stress/UPR during calcium-induced differentiation and its regulation on lysosome biogenesis/autophagy is not studied.

To connect cellular differentiation with intracellular calcium, UPR, lysosome biogenesis and autophagy, we used primary neonatal human keratinocytes (early passages less than 7) and allow them differentiate for 2-3 days using 2 mM CaCl_2_ (without starvation). Quantitative fluorescence microscopy analysis illustrated that calcium incubation increases cell length, nucelar size and the number of globular enlarged lysosomes in keratinocytes. Further, calcium-induced keratinocyte differentiation showed enhanced expression of MiT/TFE TFs similar to epidermal skin grafts and retained in the cytosol due to upregulated mTOR activity. Moreover, we found dramatic increase in autophagy flux in differentiated cells and inhibition of mTOR activity further enhanced the cellular differentiation. Biochemical approaches demonstrated that intracellular calcium levles increased during early hours of differentiaton that results in ER stress and activates ATF6 branch of UPR. Chemical inhibition of ER stress and calcium homeostasis completely abolished keratinocyte differentiation and lysosome biogenesis. Finally, our study showed that keratinocyte differentiation results in merging and dispersal of Golgi stacks and increased production of enlarged lysosomes (called Golgi-associated lysosomes, GALs); loss of Golgi function inhibited the differentiation and the concomitant lysosome biogenesis. Overall, our study provide a mechanism that extracellular calcium control the cellular fate, epidermal differentiation and skin homeostasis.

## Results

### Calcium chloride induces keratinocyte differentiation and lysosome biogenesis

Studies have shown that extracellular calcium (range from 1.0 to 1.8 mM) potentially induces the differentiation of primary human keratinocytes^10,39^. However, the mechanisms or signalling pathways behind the calcium-induced differentiation process is largely unknown. We have used commercially available primary human neonatal keratinocytes (Lonza or Invitrogen) and incubated with 2 mM CaCl2 (referred to here as +CaCl2 (2 mM) or Calcium) containing medium at cell density of around 50 – 60% confluency. Bright-field (BF) microscopy of cells showed enhanced cell-cell contacts or intercellular adhesion within 4 h (data not shown); increased cell size resembling the cellular differentiation between 24 – 48 h and the cells appeared as terminally differentiated (noticeable as tissue with transparent outgrowth on monolayer) after 9 days of CaCl_2_ incubation (Supplementary Figure 1A). Differentiated keratinocytes undergo apoptosis (rounded and floated) upon removal of calcium, suggesting that calcium-induced differentiation is an irreversible process (Supplementary Figure 1A, dedifferentiation). Moreover, keratinocyte differentiation process is specific to CaCl_2_ but not to other forms of calcium sources such as CaCO_3_ (Supplementary Figure 1A). Additionally, we observed 71±7% of keratinocytes increased their cell size (with early passaged cells) upon CaCl_2_ incubation between 48 h to 72 h (Figure 1A and Table 1)^40^. Quantitative measurement of cell length (from nucleus to cell periphery, called half-cell length, labeled as CL_H_) showed an increase in cell size by 4.7 folds in calcum-added compared to control keratinocytes (12.0±0.4 μm in Control and 56.4±1.6 μm in +CaCl_2_ cells) (Figure 1B). As expected, the size of the nuclei was increased upon incubation of keratinocytes with CaCl_2_ for 48 h to 72 h (average nuclear length: 13.4±0.2 μm in Control and 23.4±0.8 μm in +CaCl_2_ cells; average nuclear width: 10.4±0.2 μm in Control and 13.6±0.4 μm in +CaCl_2_ cells) (Supplementary Figure 1B). Consistently, the immunofluorescence (IF) intensity of cellular differentiation markers such as involucrin was dramatically increased in CaCl_2_-incubated compared to control keratinocytes (Figure 1A)^41,42^. Involucrin is a structural protein of differentiated keratinocytes that cross-link to the plasma membrane by transglutaminase I enzyme^43,44^. Upon calcium incubation, involucrin fluorescence intensity appeared as a layer beneath the cell surface compared to cytosolic diffused signal in control cells (deconvolved images in Figure 1A). In line with these results, the transcript and protein levels of involucrin were also upregulated in calcium-treated keratinocytes compared to control cell (Figure 1C). Similar to involucrin, the expression levels of other cellular differentiation specific proteins like keratin 18 and 10, but not loricrin were increased upon incubation of keratinocytes with CaCl_2_ (Figure 1C), indicating the characteristics of differentiated keratinocytes in the skin^45,46^. Thus, these studies suggest that addition of 2 mM CaCl**2** to the primary keratinocytes induces the differentiation process, resembling the epidermal differentiation^2^.

**Figure 1.**
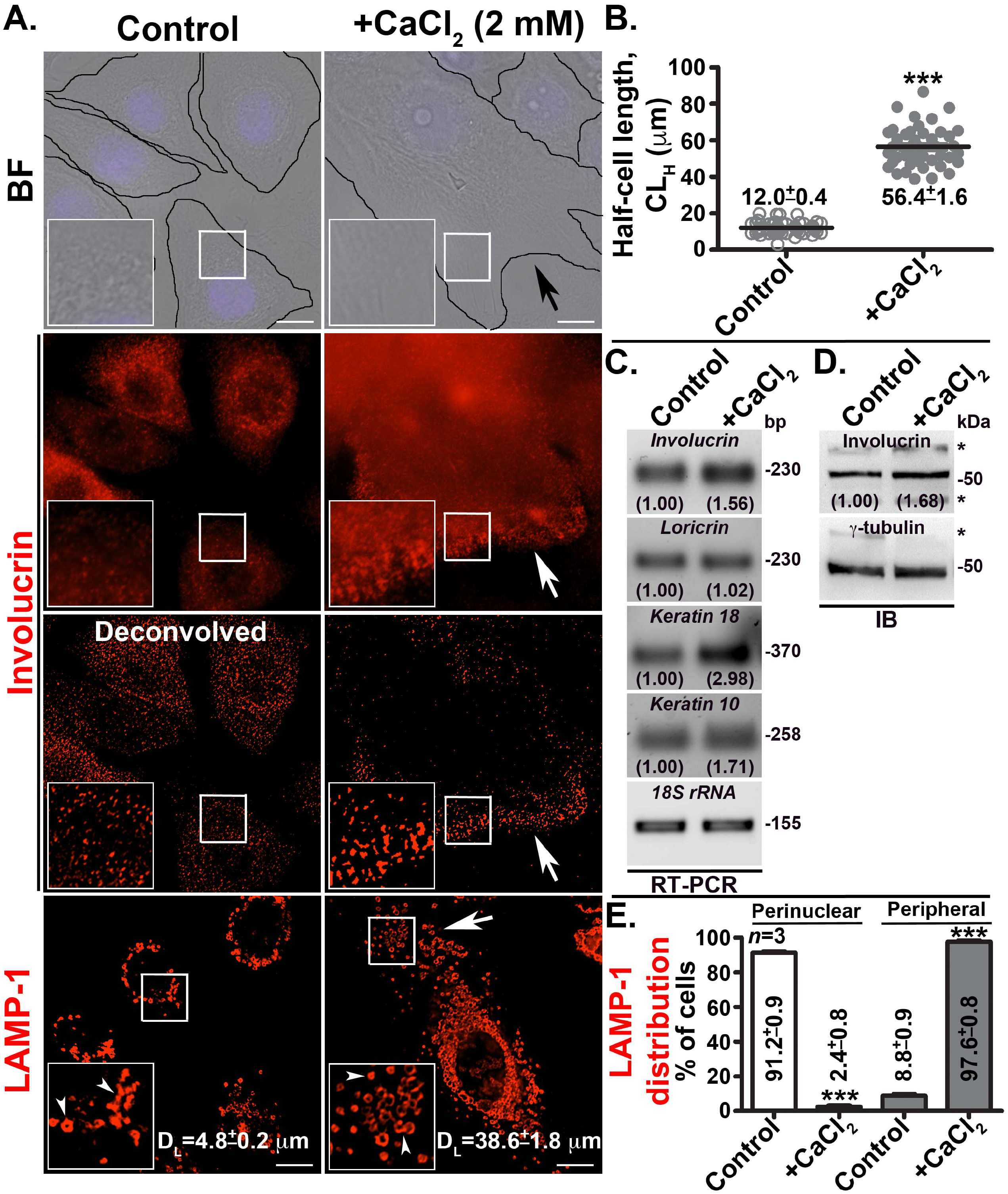
Calcium chloride incbuation induces ceulluar differentiation and increases the LAMP-1-positive compartments in human primary keratinocyte. (A) BF and IFM images of control and CaCl_2_-incubated (at 60 h) cells. Cells were immunostained for keratinocyte differentiation marker, involucrin and lysosomal protein, LAMP-1 separately. Both, un- and deconvolved images of involucrin staining were shown separately. Arrows indicate the increased cell size, cellular expression (both in cytosol and cell surface) of involucrin and LAMP-1-positive compartments. Arrowheads point to small ring-like and bigger globular lysosomes localized perinuclearly in control and distributed throughout the cytosol in differentiated cells, respectively. The insets are a magnified view of the white boxed areas. Nuclei are stained with Hoechst33258. Scale bars, 10 μm. (B) The parameter of half-cell length (in μm) was measured as the distance from nucleus to cell periphery. Approximately 60 cells (*n*=3) from each condition were measured and then plotted. Average length (mean ± s.e.m) of the cell was indicated in the graph. (C, D) Semiquantiative PCR and immunoblotting analysis of control and differentiated cells to measure the expression of various keratinocyte differentiation markers. Expected sizes (in base pairs or kilo Datons) of the transcripts and proteins were indicated. Band intensities were quantified, normalized with loading control (18S rRNA or γ-tubulin) and mentioned the fold-change on the gels. * indicates non-specific bands detected by the antibodies. (E) The distribution (perinuclear or peripheral) of LAMP-1-positive organelles in both control and differentiated cells (quantified visually). Approximately 100 cells in each condition (*n*=3) were quantified and plotted. Percentage distribution of the phenotype (mean ± s.e.m) was indicated in the graph. ***, *p*≤0.001.

**Table 1.**
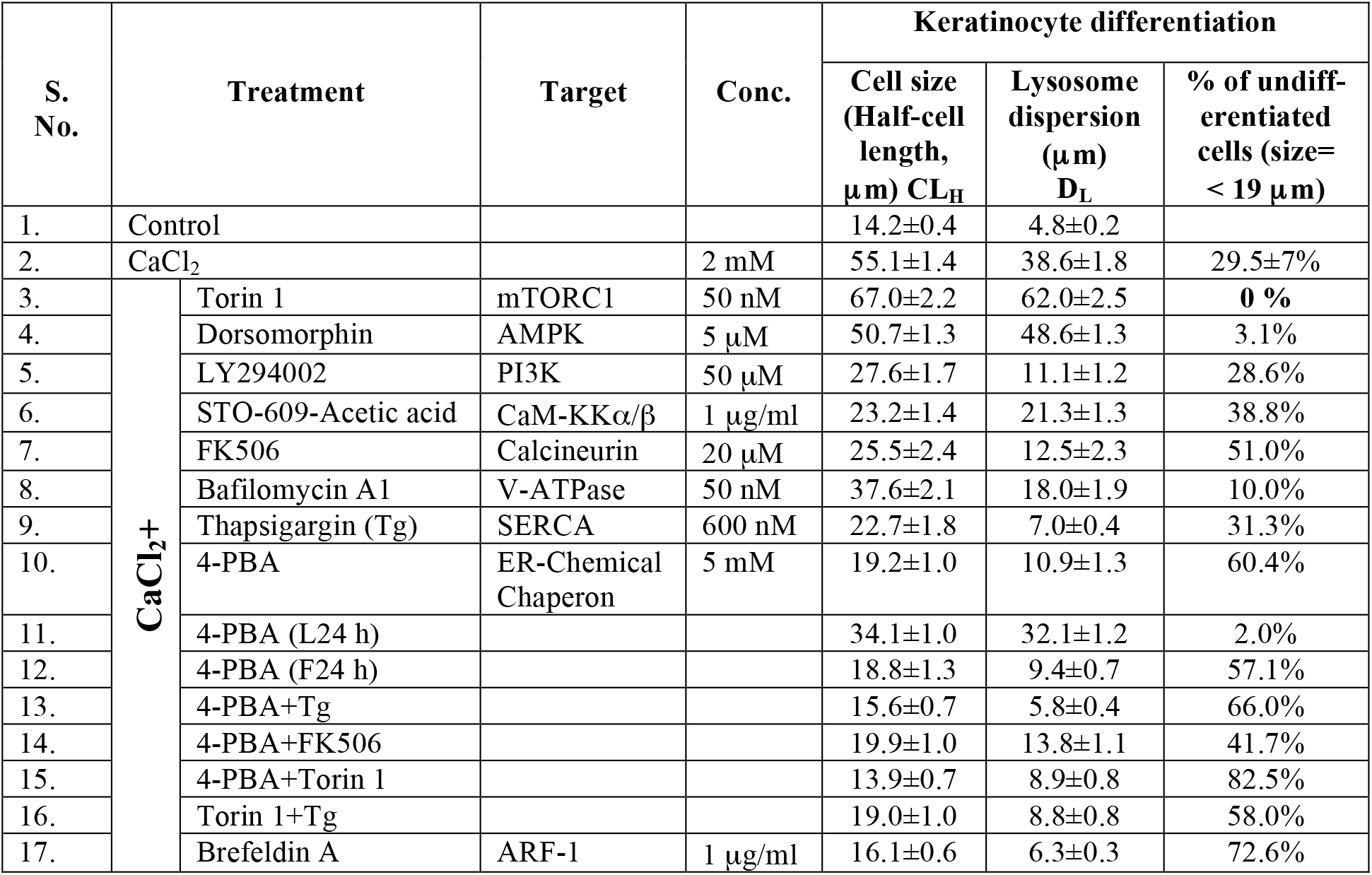
Quantification of cellular differentiation and lysosome biogenesis in keratinocytes treated with small molecule inhibitors.

We tested whether the cellular expansion during keratinocyte differentiation alters the number and size of intracellular organelles. Immunofluorescence microscopy (IFM) analysis of differentiated keratinocytes showed dramatic increase in size and number of LAMP-1-positive compartments (corresponds to lysosomes, see below) compared to EEA1-positive early endosomes or STX13-positive recycling endosomes (Figure 1A and Supplementary Figure 1C). Moreover, these organelles were majorly distributed to the cell periphery in differentiated cells compared to control keratinocytes (Figure 1A and 1E), suggesting that the differentiation also alters the position and number of LAMP-1-positive intracellular organelles. Quantitative measurement of lysosome distribution in the cells (from nucleus to cell periphery, called lysosome dispersion, labeled as D_L_) showed 8 folds increase in calcium-added compared to control keratinocytes (4.8±0.2 μm in control and 38.6±1.8 μm in +CaCl_2_ cells) (Table 1 and Figure 1A). We used this parameter as well as cell size to distinguish the differentiated keratinocytes (see below). Quantitative IFM analysis showed that the LAMP-1-positive organelles in CaCl_2_-treated cells are acidic as stained by lysotracker (Pearson’s coefficient, r=0.16±0.02 in Control and 0.48±0.02 in +CaCl_2_ cells) and proteolytically active as judgeed by increased fluorescence intenstity of DQ-BSA (r=0.20±0.02 in Control and 0.44±0.02 in +CaCl_2_ cells) (Figure 2A). As expected, these organelles were majorly positive for known lysosome-associated proteins such as Arl8b-GFP^47^ and Rab9^48^ compared to late endosomal protein HA-Rab7A^49^ (Figure 2A and Supplementary Figure 1C), confirming the characteristics of lysosomes. Consistently, the lysosomal enzyme activity (such as Glucocerebrosidase, GBA) as measured biochemically was enhanced by 2.15 fold in differentiated cells compared to control keratinocytes (2.15±0.12 folds in +CaCl_2_ compared to control cells) (Figure 2B). Likewise, the expression levels (but not transcript levels, see below) of lysosome encoded hydrolases (GBA) and the membrane localized (LAMP-1 and -2) or associated (Rab9) proteins but not hydrolase transporters (LIMPII) were significantly increased in differentiated cells compared to control keratinocytes (Figure 2C). However, the cell surface expression of LAMP-1 and -2 were moderately increased in CaCl_2_-treated compared to control cells (1.46±0.13 folds for LAMP-1 and 1.50±0.20 folds for LAMP-2 in +CaCl_2_ cells) (Figure 1D). Overall, these studies indicate that the extracellular CaCl_2_ induces lysosome biogenesis along with keratinocyte differentiation.

**Figure 2.**
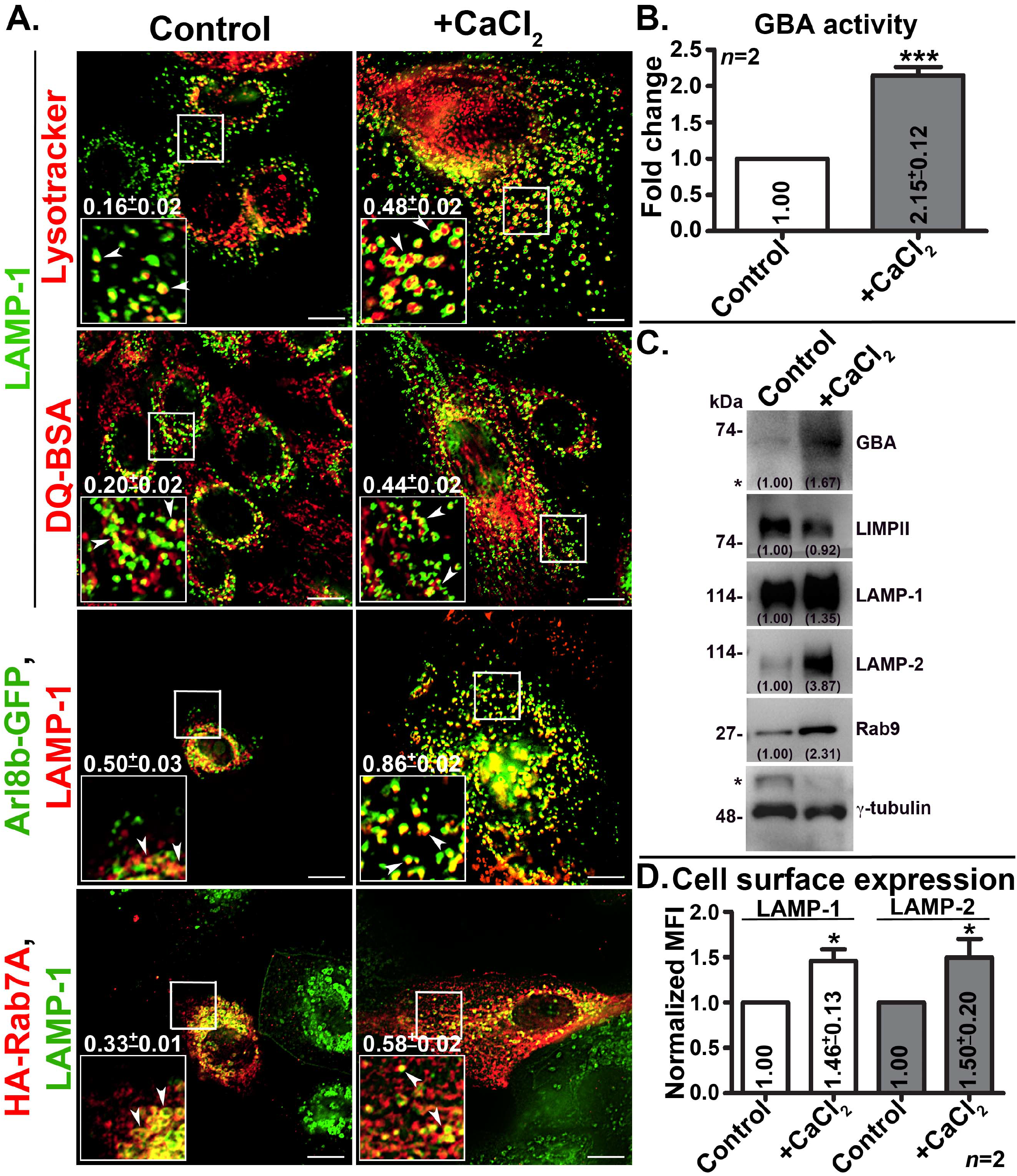
Differentiated keratinocytes contain active globular/enlarged lysosomes. IFM analysis of control and differentiated kerationocytes that were either internalized with lysotracker or DQ-BSA or transfected with Arl8b-GFP or HA-Rab7A plasmids, and then fixed, stained with LAMP-1 and imaged. Arrowheads point to the LAMP-1-positive organelles with respect to other lysosomal markers. The degree of colocalization between the two markers was measured as Pearson’s coefficient value ‘*r*’, indicated separately (mean±s.e.m.). The insets are a magnified view of the white boxed areas. Scale bars, 10 μm. (B) Biochemical anlysis of Glucocerebrosidase (GBA) activity to represent the lysosome enzyme activity in control and differentiated cells. Activity values were normalized with cell number and plotted the average fold change in activity (mean ± s.e.m) as indicated (*n*=2). ***, *p*≤0.001. (C) Immunoblotting analysis of control and differentiated cells to measure the expression of lysosome carrier/resident or its associated proteins. Band intensities were quantified, normalized with loading control (γ-tubulin) and mentioned the fold-change on the gels. * indicates non-specific bands detected by the antibodies. (D) Cell surface levels of LAMP-1 and -2 in control and differentiated cells measured using flow cytometry. Fold change in mean fluoresence intensity (MFI) was calculated (from two independent experiments with quadruplicates each) and then plotted (mean±s.e.m.). *, *p*≤0.05.

### Increased lysosome biogenesis during keratinocyte differentiation is independent of the pathway involving MiT/TFE transcription factors

Recent studies have reported that lysosome biogenesis is under the control of MiT/TFE TFs involving TFEB (transcript factor EB), TFE3 (transcription factor E3) and MITF (microphthalmia-associated transcription factor)^26,28,50^. We investigated whether these TFs are involved in keratinocyte differentiation and concomitantly upregulate the lysosome biogenesis. Histocytochemistry analysis of epidermal skin grafts showed enhanced TFEB fluorescence intensity in the involucrin-positive granular layer, positioned between basal layer and cornified layer (stratum cornium) of epidermis (Figre 3A). Moreover, the TFEB-positive layer showed lower keratin14 expression (Supplemnetary Figure 1D), a marker for epidermis basal layer (corresponds to stratum spinosum), suggesting that TFEB expression is upregulated in the stratum graulosum layer. Immunoblotting analysis of differentiated cells displayed increased protein levels compared to control cells although the transcript levels of MiT/TFE TFs were unchanged (Figure 3B and Supplementary Figure 1E). However, IFM and nuclear fractionation analysis showed the nuclear localization of GFP-TFEB or the endogenous TFE3 or MITF was not increased in Calcium-treated compared to control cells (Figure 3C ad 3D). Consistently, the expression of several target genes (*HPRT1, LAMP1, MCOLN1, NAGLU1, ATP6, CLCN7, CTSD* and *GLA*) associated with the lysosome biogenesis under the control of MiT/TFE TFs was not altered in differentiated cells compared to control keratinocytes (Supplementary Figure 1F). These results suggest that the enhanced expression of MiT/TFE TFs during calcium-induced keratinocyte differentiation may not contribute to the observed upregulated lysosome biogenesis.

**Figure 3.**
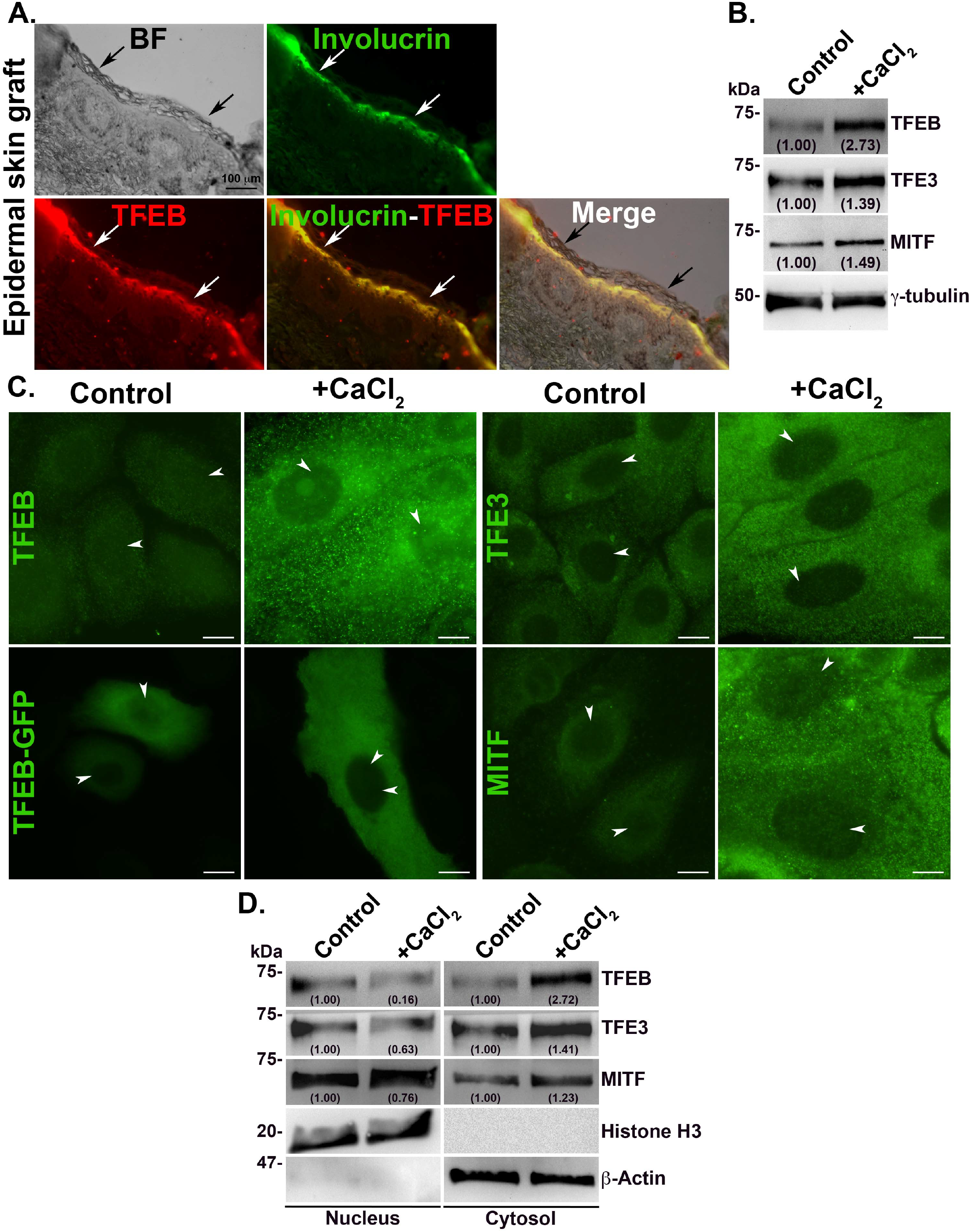
Differentiated keratinocytes of human skin or primary cells increases the expression of MiT/TFE TFs and retained in the cytosol. (A) BF and IFM of epidermal skin graft that was immunostained for keratinocyte differentiation maker involucrin and TF TFEB. Black arrows point to the cornified outermost layer located above the immunostained keratinocyte layer. White arrows indicate the differentiated keratinocytes (positive for involucrin) that showed enhanced TFEB expression. Scale bars, 100 μm. (B) Immunoblotting analysis of cells for the expression levels of MiT/TFE TFs (TFEB, TFE3 and MITF). Band intensities were quantified, normalized with loading control (γ-tubulin) and mentioned the fold-change on the gels. (C, D) IFM and nuclear fractionation analysis of keratinocytes for the localization of endogeneous MiT/TFE TFs. Cells transfected with ectopically expressed TFEB-GFP was shown separately. Arrowheads point to the localization of TFs to the nucleus. Scale bars, 10 μm. Fractionated nucleus and cytosolic pools were probed with indicated antibodies. Band intensities were quantified, normalized with respective loading controls (Histone H3 for nuclear and β-Actin for cytosolic fraction) and mentioned the fold-change on the gels.

### Calcium-dependent keratinocyte differentiation upregulates the autophagy flux and lysosome biogenesis independent of mTOR activity

As reported, MiT/TFE TFs also regulate the expression of genes that control the autophagy, a cellular clearence pathway in all cell types^26,51–53^. Moreover, mTORC1 has been shown to phosphorylate the MiT/TFE TFs and retain them to the cytosol^28^; and calcineurin dephosphorylates these TFs depending on the MCOLN1-mediated calcium release from lysosomes^30^. Additionally, a recent study has shown that mTORC complexes (1 and 2) regulate the formation of functional epidermis *in vivo*^20^. We investigated whether keratinocyte differentiation alters the mTOR-dependent autophgy process. Immunoblotting analysis showed calcium addition to the keratinocytes upregulates the activity of mTOR and the related autophagy pathway components (Figure 4A). IFM analysis revealed that enhanced phospho-mTOR (active mTOR or pmTOR) fluorescence intensity corresponds to its protein levels in the differentiated keratinocytes compared to control cells (Figure 4B). Interesingly, the localization of pmTOR to the lysosome membranes^29^ was increased in differentiated kerationcytes compared to control cells (r=0.61±0.05 in Control and 0.91±0.03 in +CaCl_2_ cells) (Figure 4B), indicating that pmTOR possibly enhance the phosphorylation of MiT/TFE TFs that lead to their retention to the cytosol (Figure 3C). Studies have shown that mTORC2 phosphorylates AKT for its activation, which further phosphorylates mTORC1^54^. As expected, the phosphorylation of AKT was significantly increased during keratinocyte differentiation, which is in line with the activated mTORC1 in the cells (Figure 4A). Chemical inhibition of mTORC1 using Torin 1 enhanced the differentiation of keratinocytes to 100% (Table 1) and the cells possess deformed nucleus indicative of initiation of nucleophagy (arrow in Figure 4C). Surprisingly, the TFEB TF majorly localized to the lysosome membranes upon treatment with Torin 1 in calcium-treated compared to control keratinocytes (Figure 4C). These results indicate that calcium-induced differentiation activates mTORC1 and its inactivation by active-site inhibitor, sequesters the MiT/TFE TFs such as TFEB onto the lysosome membranes.

**Figure 4.**
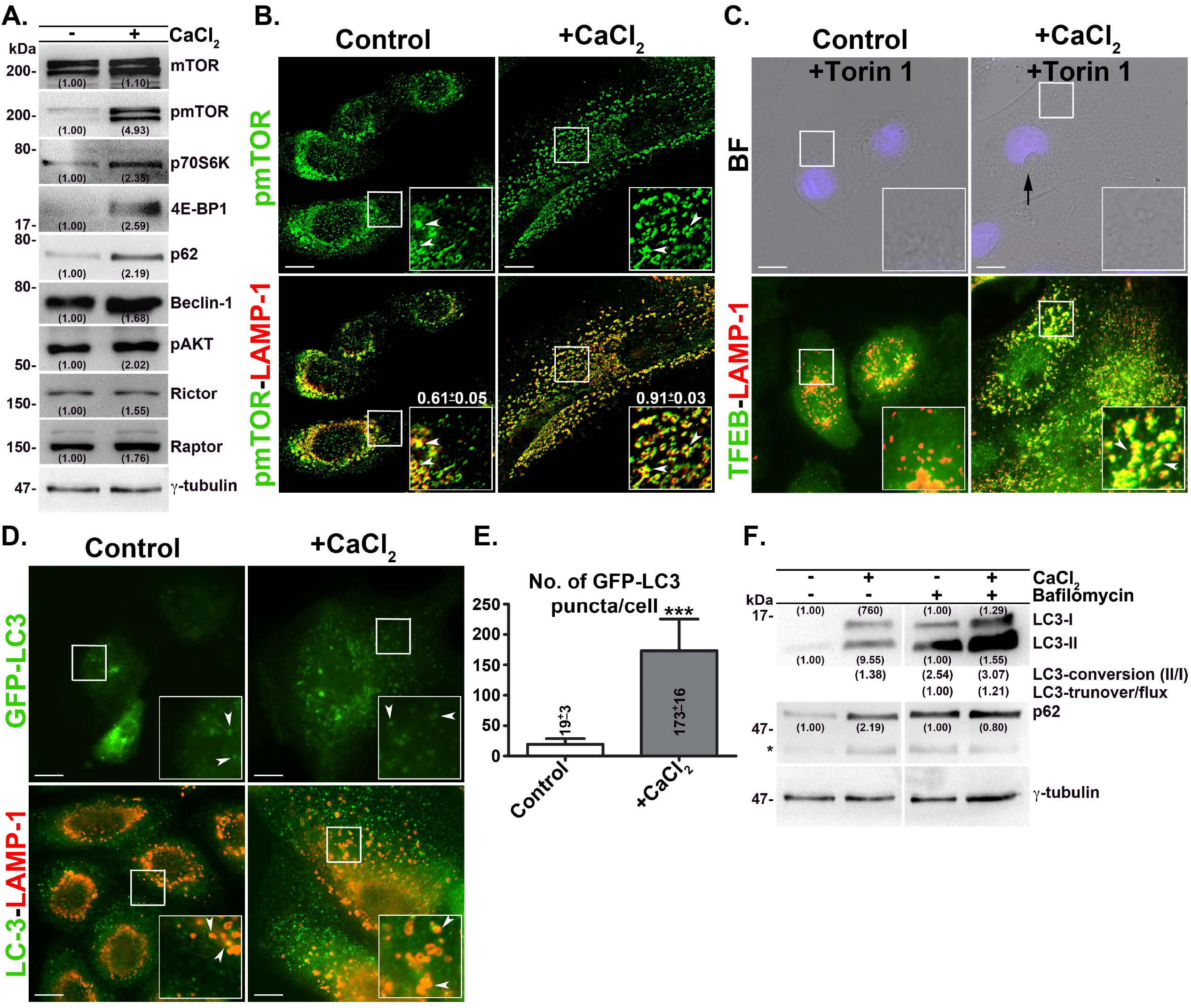
Calcium-induced keratinocyte differentiation activates the mTOR and autophagy flux; and chemical inhibition of mTOR activity enhances the differentiation of keratinocytes. (A) Immunoblotting analysis of multiple autophagy specific genes, which indicate the status of autophagy process in the keratinocytes. Phosphorylation status of few proteins indicated separately. Band intensities were quantified, normalized with loading control (γ-tubulin) and mentioned the fold-change on the gels. (B-D) BF and IFM analysis of keratinocytes for the locazation of mTOR, TFEB or LC3 with respective to LAMP-1 or GFP-LC3 alone. Cell were treated with mTOR inhibitor Torin 1 (50 nM) in C. Arrowheads point to the LAMP-1-positive organelles with respect to other proteins or GFP-LC3-positive autophagosomes (in D). Black arrow in C indicates the deformed nucleus. The degree of colocalization between the two markers was measured as Pearson’s coefficient value ‘r’, indicated separately (mean±s.e.m.). Nuclei are stained with Hoechst33258. The insets are a magnified view of the white boxed areas. Scale bars, 10 μm. (E) Visual quantification of number of autophagosomes (positive for GFP-LC3) in control and differentiated cells. Average number of puncta from 10 cells was quantified, plotted and indicated (mean ± s.e.m.). ***, p<0.001. (F) Quantification of autophagy flux by immunoblotting in cells as indicated. Cells were treated with or without bafilomycin A1 during the differentiation process. Band intensities were quantified, normalized with loading control (γ-tubulin) and mentioned the fold-change on the gels. The rate of LC3 conversion (LC3-II/LC3-I) and autophagy flux/turnover (fold change in LC3 conversion with bafilomycin A1) were indicated separtely.

Next, we studied the dynamics of autophagosome formation by monitoring the localization and conversion of LC3-I into II. IFM analysis showed increased punctate localization of GFP-LC3 in differentiated cells compared to the control cells (Figure 4D). As expected, visual quantification GFP-LC3 puncta (indicative of autophagosomes) revealed that the number is upregulated by 9 folds in +CaCl_2_ cells compared to control cells (Figure 4E). Moreover, colocalization of endogenous LC3 with lysosomes (marked by LAMP-1) was moderately increased in differentiated keratinocytes compared to control cells (Figure 4D) in addition to the increased LC3 expression (Figure 4F), indicating slightly altered autophagy flux in these cells. Quantitative measurement of LC3 conversion from I to II or the net autophagy flux measued using bafilomycin A1 (inhibitor of V-ATPase, blocks the fusion of autophagosome with lysosomes^55^) was dramatically increased in calcium-induced differentiated compared to control cells (Figure 4F). It has been shown that autophagy substrate capture receptor p62 (also called SQSTM1) facilitates degradation of autophagosome membranes by binding to LC3-II^56^. Immunoblotting analysis showed significant increase in the protein levels of p62 upon incubation of keratinocytes with CaCl_2_, however, those levels were slightly reduced upon bafilomycin A1 treatment (Figure 4F and also see bafilomycin-treatment in Figure 5A). Overall, these results suggest that the enhanced autophagy during keratinocyte differentiation is independent of mTOR activity. Nevertheless, calcium addition to the keratinocytes possibly enhances the autophagy flux, but slightly delayed autophagosome fusion with lysosomes. Altogether, these data suggest that mTOR activity is dispensable during keratinocyte differentiation.

**Figure 5.**
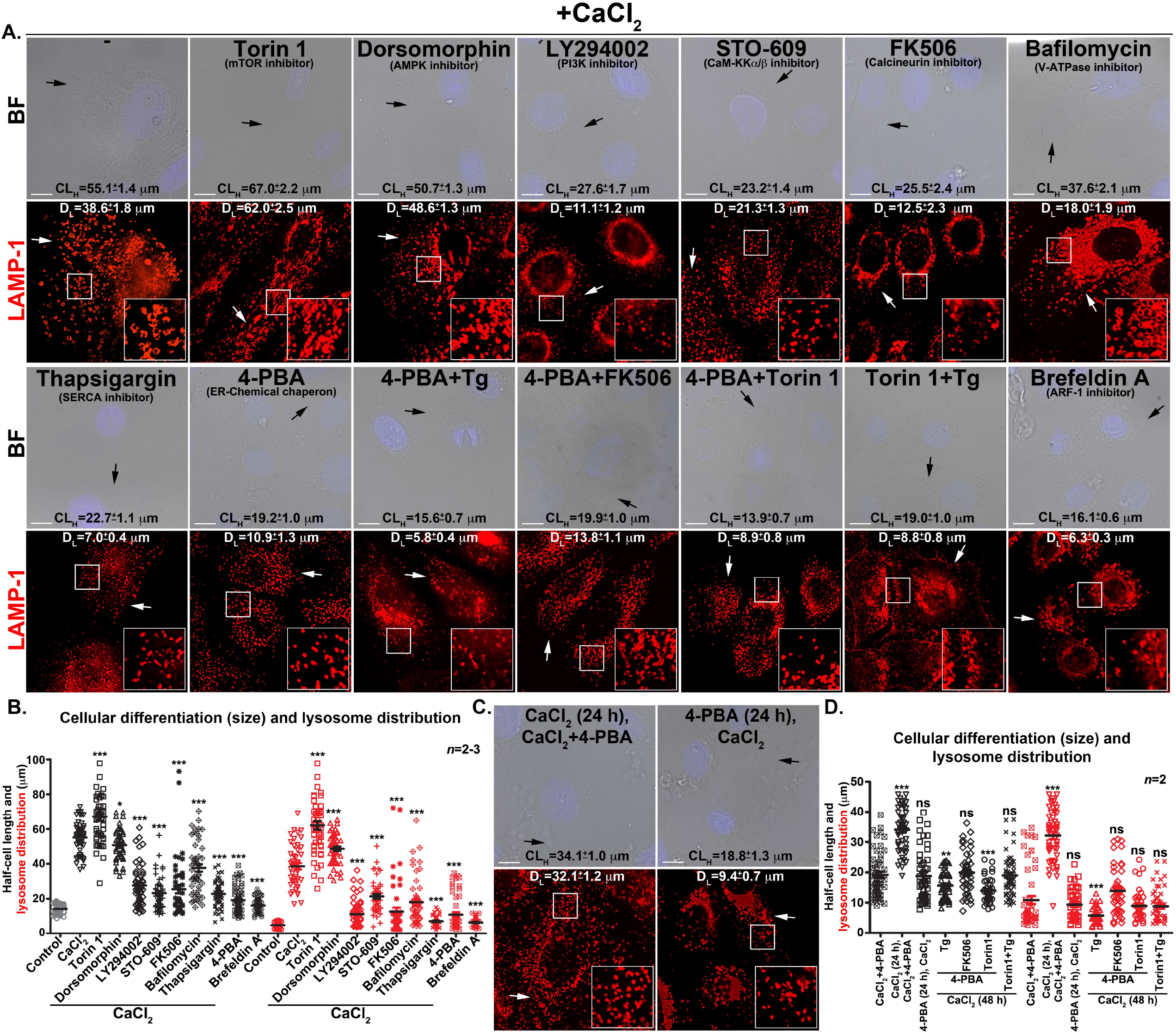
ER stress and Golgi function regulates the keratinocyte differentiation and lysosome biogenesis. (A, C) BF and IFM images of keratinocytes representing each condition and were used for measuring cell size and lysosome biogenesis (number and distribution). Cells were co-treated with the indicated compounds (concentrations were listed in Table 1) and CaCl_2_ for 48 h (entire duration of differentiation), fixed and stained for LAMP-1. In C, cells were pre- or post-treated with 4-PBA along with CaCl_2_ as described. Black arrows points to the cell size and white arrows indicate the distribution/morphology of lysosomes. Nuclei are stained with Hoechst33258. The insets are a magnified view of the white boxed areas. Scale bars, 10 μm. (B, D) Quantification of cellular differentiation and lysosome biogenesis were measured as half-cell length (CL_H_, black symbols) and lysosome dispersion (DL, red symbols), respectively in the indicated treatments. Both CL_H_ and DL were quantified as μm from the nuclues towards cell surface (approximately 60-80 cells, *n*=2-3) and then plotted. Average CL_H_ and D_L_ (in μm) for each treatment were indicated on IFM images and also in Table 1 (mean ± s.e.m.). Statstical analysis between drug treatment and CaCl_2_-alone was indicated. *, *p*≤0.05; **, *p*≤0.01; ***, *p*≤0.001 and ns, not significant.

### ER stress regulates keratinocyte differentiation and lysosome biogenesis

To understand the mechanism of calcium-induced keratinocyte differentiation and its molecular link with lysosome biogenesis, we evaluated the role of several components of calcium-dependent signalling pathways including lysosome/Golgi biogenesis by using specific chemical inhibitors or modulators (Table 1). Here, we have used the cell size (CL_H_) and lysosome dispersion (D_L_) as parameters for quantifying the status of keratinocyte differentiation and lysosome biogenesis, respectively. Studies have been shown that cytosolic calcium either sensed by calmodulin (proximal calcium sensor), which then activates calmodulin kinase and calcineurin or lead to endoplasmic reticulum (ER) stress. Moreover, these processes independently regulate the lysosome biogenesis/autophagy by non-canonical mTOR pathway^35,57^. Additionally, AMPK downstream signalling molecule PI3K has been shown to play critical role in keratinocyte differentiation^21,22^. Chemical inhibition of calmodulin KKα/β (STO-609-acetic acid) or calcineurin (FK506) moderately reduced (by 0.42 and 0.46 folds, respectively) the calcium-dependent keratinocyte cell size/differentiation, but the calcineurin inhibition showed significantly reduced lysosome biogenesis compared to CaCl_2_ alone or in combination with STO-609-acetic acid (Figure 5A, 5B and Table 1). The calcineurin inhibition effect on lysosome biogenesis was unexpected, since this phosphatase is known to dephosphorylate and translocate the TFs of lysosome biogenesis^30^. We hypothesize that the reduction in calcium-induced keratinocyte differentiation (0.46 folds)/lysosome biogenesis (0.32 folds) upon calcineurin inhibition is very likely due to decreased dephosphorylation of ER chaperon calnexin during ER stress^58^. In contrast, AMPK (dorsomorphin) inhibition showed no major change in calcium-dependnet keratinocyte differentiation/lysosome biogenesis compared to CaCl_2_-incubated cells (Figure 5A, 5B and Table 1). However, PI3K (LY294002) or V-ATPase (bafilomycin A1) inhibition showed modest reduction in calcium-induced keratinocyte differentiation (0.5 folds for PI3K and 0.68 folds for V-ATPase) and lysosome biogenesis (0.29 folds for PI3K and 0.47 folds for V-ATPase) compared to CaCl_2_ alone-treated cells (Figure 5A, 5B and Table 1). As expected, mTORC1 inhibition (Torin 1) showed accelerated differentiation/lysosome biogenesis compared to CaCl_2_-incubated cells (Figure 5A, 5B and Table 1), suggesting mTOR function is dispensible during calcium-mediated keratinocyte differentiation. Cotreatment of keratinocytes with ER stress modulator such as thapsigargin (Tg, an inhibitor of SERCA-calcium pump and blocks the influx of calcium into ER) and CaCl_2_ significantly reduced the differentiation (0.41 folds) and completely abolished the lysosome biogenesis (0.18 folds) compared to CaCl_2_ alone (Figure 5A, 5B and Table 1; see below). However, ER stress attenuator 4-PBA (4-Phenylbutyric acid, a chemical chaperon that attenuates ER stress by increasing protein folding)^59^ did not enhance either the calcium-dependent keratinocyte differentiation (0.35 folds) nor lysosome biogenesis (0.28 folds) compared to CaCl_2_ alone (Figure 5A, 5B and Table 1). To understand the role of ER stress in keratinocyte differentiation and its concomittent lysosome biogenesis, we incubated the cells with CaCl_2_ for 24 h and then co-incubated with 4-PBA for another 24 h or pretreated the cells with 4-PBA alone and then incubated with CaCl_2_ (Figure 5C, 5D and Table 1). Pretreatment of cells with 4-PBA inhibited both the keratinocyte differentiation and lysosome biogenesis equivalent to that of 4-PBA and CaCl_2_ co-treated cells (Table 1). In contrast, post treatment of 4-PBA did not show any strong effect on CaCl_2_-induced keratinocyte differentiation/lysosome biogenesis (Figure 5C, 5D and Table 1). These results suggest that (1) extracellular calcium very likely generates ER stress that possibly regulates the cell size and lysosome biogenesis; and (2) an initial calcium induced cellular signalling or ER stress is sufficient to differentiate the primary human keratinocytes. Studies have shown that 4-PBA attenuates Tg-induced ER stress in tumor microenvironment^60^. We tested whether 4-PBA can rescue the Tg-dependent inhibition of calcium-mediated keratinocyte differentiation (Figure 5A, 5D and Table 1). Co-treament of keratinocytes with Tg and 4-PBA with CaCl_2_ showed significant reduction in cellular size (differentiation) as well as lysosome biogenesis compared to Tg+CaCl_2_ or 4-PBA+CaCl_2_-incubated cells (Figure 5A, 5D and Table 1). This result suggests that 4-PBA-mediated ER stress attenuation is not sufficient to rescue the blockade in cellular differentiation observed with SERCA-pump inhibitor Tg. Additionally, coincubation of keratinocytes with 4-PBA, FK506 and CaCl_2_ showed similar level of differentiation/lysosome biogenesis as observed in cells treated with 4-PBA+CaCl_2_ or FK506+CaCl2 (Figure 5A, 5D and Table 1). Moreover, Torin 1-mediated enhanced keratinocyte differentiation (including lysosome biogenesis) was drastically reduced upon coincubation with 4-PBA or Tg along with CaCl_2_ (Figure 5A, 5D and Table 1). These results indicate that (1) attenuation of ER stress at early hours by using 4-PBA or an additional ER stress caused by Tg blocks keratinocyte differentiation; (2) an optimized ER stress resulted by extracellular high CaCl_2_ (see below) is sufficient to induce the keratinocyte differentiation and any added ER stress possibly block the differentiation; and (3) artificial nutritional stress as possibly being created by mTORC1 inhibition, is the only additive factor to the keratinocyte differentiation and it acts in an ER stress-dependent manner. Additionally, we speculate that ER stress and calcium refilling through SERCA pump are likely regulate both keratinocyte differentiation and lysosome biogenesis. Overall, these studies indicate that cellular calcium levels and ER stress may contribute to the skin epidermal differentiation.

### Cytosolic calcium activates UPR TFs and modulates lysosome biogenesis during keratinocyte differentiation

We evaluated the role of cytosolic calcium [Ca^2+^] in the process of keratinocyte differentiation and enhanced lysosome biogenesis. Previous studies speculated that extracellular calcium elevates and activates intracelluar calcium signalling that lead to keratinocyte differentiation^1,13^. Endoplasmic reticulum and its calcium pumps IP3R (inositol triphosphate receptor), RYR (ryanodine receptor) and SERCA (sarco endoplasmic reticulum calcium ATPase) maintain the intracellular calcium concentration/gradent either by releasing calcium to the cytosol (by IP3R, RYR) or by pumping extra calcium into ER (by SERCA). However, the critical role of cytosolic calcium on the MiT/TFE TFs activity or its connection to unfolded protein response (UPR) during keratinocyte differentiation is unknown. Kinetic measurement of intracellular calcium using Fluo-4 NW revealed that the calcium levels were significantly higer in the initial two hours of keratinocyte differentiation and restored to or reduced than the basal calcium levels at 48 h (Figure 6Ai), suggesting a role for early calcium flux that may lead to the calcium-dependent endoplasmic reticulum (ER) stress. Moreover, this data suggest that differentiated keratinocytes maintain lower calcium concentration than control cells (Figure 6Ai), inspite of constant exposure of cells to the high concentration of extra cellular calcium. As predicted, the peak calcium flux observed in differentiated keratinocytes was abolished and appeared as equivalet to that of the control cells upon treatment with Tg for 6 h (blocks the influx of calcium into ER through SERCA^61^) (Figure 6Aii). Consistently, calcium chelator EGTA had no effect (at 6 h) on intracellular calcium flux in the keratinocytes with or without addition of extracellular calcium (Figure 6Aii). Studies have shown that calcium release from the ER modulates the activity of mTOR and its downstream TFEB/TFE3-mediated lysosomal gene expression^35^. Interestingly, treatment of keratinocytes with mTOR inhibitor Torin 1 (for 6 h) moderately increased the cytosolic calcium both in Torin 1-alone and CaCl_2_+Torin 1-incubated cells compared to only CaCl_2_-treated cells (Figure 6Ai and 6Aii). As expected, intracellular calcium level was lower than the basal level (slighly higher than the CaCl_2_-alone) upon treatment of keratinocytes with CaCl_2_+Torin 1 for 48 h (Figure 6Aiii). Thus, these results explain the enhanced keratinocyte differentiation observed with CaCl_2_+Torin 1 compared to CaCl_2_ alone-incubated cells (Table 1). Moreover, these studies suggest that the observed calcium peak at 2 h upon calcium addition to the keratinocytes is possibly due to increased calcium release from the ER and later calcium re-enters the ER through SERCA pump. We hypothesize that this process possibly maintains the lower cytosolic calcium concentration during keratinocyte differentiation.

**Figure 6.**
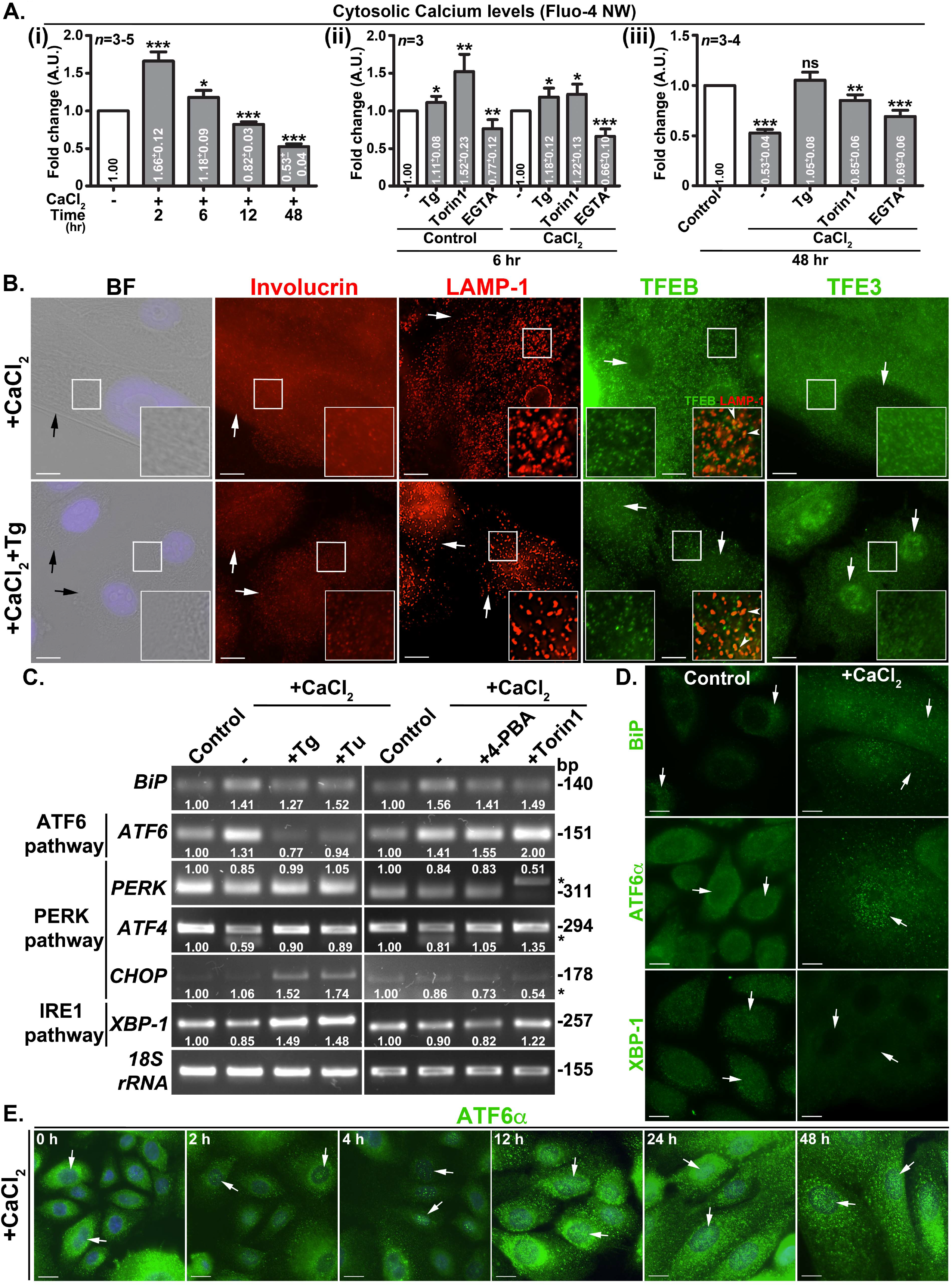
Differentiation of keratinocytes elevates cytosolic calcium at early phase and activates UPR TF ATF6α. (A) Measurement of intracellular calcium using Fluo-4 NW in keratinocytes at different time points of differentiation. In (i), cells were treated with CaCl_2_ for respective time points. In (ii and iii), cells were incubated with the indicated compounds alone or coincubated with CaCl_2_ for the respective time points. In all conditions, cells were incubated with Fluo-4 NW for 2-3 hours either prior to or during the treatment condition. The fluorescence intensities were measured, normalized with cell numbers and plotted the fold changes with their respective controls (mean ± s.e.m.). Note that EGTA was used as the negative control in the experiments. *, *p*≤0.05; **, *p*≤0.01; ***, *p*≤0.001 and ns, not significant. (B, D, E) BF and IFM analysis of keratinocytes that were incubated with CaCl_2_ alone or CaCl_2_ with Tg for 48 h. Cells were fixed, immunostained and imaged. Black arrows point to the cell size. White arrows indicate the expression of proteins or distribution of lysosomes or localization of respective TFs to the nucleus. Arrowheads point to the localization of TFEB to the lysosomes. Nuclei are stained with Hoechst33258. The insets are a magnified view of the white boxed areas. Scale bars, 10 μm. (C) Transcript analysis of ER stress genes in the keratinocytes that were treated with either CaCl_2_ alone or CaCl_2_ with the indicated compounds for 48 h. Expected size (in base pairs) of each transcript was indicated. Band intensities were quantified, normalized with loading control (18S rRNA) and mentioned the fold change on the gels. * indicates non-specific bands amplified during the PCR.

We tested whether the inhibition of SERCA function alters the differentiation of kertinocytes. IFM analysis of Tg-treated calcium-incubated keratinocytes showed reduced involucrin expression and cell size compared to differentiated cells (Figure 6B and Table 1). Consistently, LAMP-1-positive lysosomes were reduced in CaCl_2_-treated cells upon coincubation with Tg (Figure 6B). As expected, the localization of TFEB to the nucleus was not altered (arrows), but its localization to the lysosomes was moderately reduced (arrowheads) upon co-treatment of keratinocytes with Tg and CaCl_2_ (Figure 6B, inset of TFEB panel). In constrast, the nuclear localization of TFE3 was enhanced upon treatment of keratinocytes with Tg and CaCl_2_ (Figure 6B), however, it did not contribute to the increased lysosome biogenesis or keratinocyte differentiation (Table 1 and Supplementry Figure 1). Earlier studies have shown that increased cytosolic calcium upregulates ER proteostasis pathways such as unfolded protein response (UPR)^35^. We evaluated whether ER stress is enhanced during keratinocyte differentiation process. Transcript analysis of keratinocyte lysates showed moderate increase in ER chaperon BiP in CaCl_2_-cells compared to control cells (Figure 6C). Interestingly, increased BiP transcript levels in the differentiated cells were not further enhanced with ER stress inducers Tg or Tu (tunicamycin), nor reduced with ER stress attenuator 4-PBA or mTOR inhibitor Torin 1 (Figure 6C). Consistently, immunoblotting and IFM analysis showed notable increase in protein levels of BiP and another chaperon calnexin in the differentiated cells compared to control cells (Figure 6D and Supplementary Figure 2A, IFM data not shown for Calnexin), indicating a moderate upregulation of ER stress in the differentiated keratinocytes.

Next, we examined the status of three branches of UPR (ATF6, PERK and IRE1) pathway during calcium-induced keratinocyte differentiation. Transcript analysis showed an increased transcript levels of ATF6 but not PERK (PERK, ATF4, CHOP)-or IRE1 (XBP-1)-dependent pathway genes during calcium-dependent keratinocyte differentiation (Figure 6C). Interestingly, the upregulation of ATF6 transcripts upon calcium-treatment was similar even after the incubation of cells with 4-PBA or Torin 1 (Figure 6C). However, the treatment of calcium-incubated cells with 4-PBA or Torin 1 did not alter the transcription profile of PERK or IRE1-pathway genes (Figure 6C). In contrast, Tg or Tu treatment during keratinocyte differentiation significantly reduced the transcript levels of ATF6 (Figure 6C), consistent with a block in keratinocyte differentiation (Figure 5A, 5B and Table 1, data not shown for tunicamycin). Additionally, moderately enhanced CHOP (PARK-pathway) and XBP-1 (IRE1 pathway), but not PERK or ATF4 transcripts were observed upon treatment with Tg or Tu during differentiation (Figure 6C). It has been shown that ATF6 traffics from ER to Golgi during ER stress and undergo cleavage by S1P and S2P proteases that results in release of cytosolic ATF6 TF^62^. Moreover, studies have also shown that activated ATF6 TF regulates the lysosome biogenesis and autophagy^63^. Immunoblotting analysis showed marginally increased ATF6 TF and PERK protein levels compared to IRE1α or spliced XBP-1 in keratinocyte differentiated cells (Supplementary Figure 2A). Interestingly, the nuclear localization of ATF6 but not XBP-1 was increased in differentiated keratinocytes compared to control cells (Figure 6D). As expected, time kinetics studies showed enhanced localization of ATF6 to the nucleus between 12 h to 48 h of calcium-incubated keratinocytes (Figure 6E). Thus, these studies suggest that ATF6 possibly play a key role in calcium-dependent keratinocyte differentiation.

### ER stress dependent lysosomes are derived from Golgi

To investigate the origin of ER stress-dependent lysosomes, we evaluated the role of other oraganelles in the differentiated keratinocytes. IFM analysis showed increased fluorescence intensity of calnexin (ER resident protein, not shown), consistent with the upregulated proteins levels (Supplemental Figure 2A) in differentiated compared to control cells. Moreover, cellular distribution of early, recycling and late endosomes was not affected upon keratinocyte differentiation (Supplemental Figure 1C). However, the localization of late endosomal/lysosomal protein Rab9 to LAMP-1-positive lysosomes near the perinuclear area was increased in calcium-treated compared to control keratinocytes (Supplemental Figure 1C). Interestingly, the dispersal and fragmentation of both *cis*- and *trans*-Golgi (labeled with Golgi-tethering proteins GM130 and Golgin 97/p230, respectively), but not transitional ER (marked by ERGIC-53) was dramatically increased in differentiated keratinocytes compared to control cells (Figure 7A and Supplemental Figure 2B-2F). To our surprize, the fragmented *cis*- and trans-Golgi (but not the transitional ER) were positive for LAMP-1-positive lysosomes in differentiated cells compared to control cells (Figure 7A and Supplementary Figure 2B), indicating that these organelles possibly originated from Golgi compartments. Moreover, keratinocyte differentiation showed increased colocalization of GM130 with p230 suggesting a merge between the Golgi compartments (Figure 7A and Supplementary Figure 2B). Consitently, the inhibition of Golgi function using brefeldin A (inhibits ARF-1 activity) completely abolished the cellular differentiation and lysosome biogenesis in keratinocytes (Figure 5A, 5B and Table 1), suggesting that Golgi regulates lysosome biogenesis. Due to association of Golgi-tethering factors with these lysosomes, we named these organelles as Golgi-associated lysosomes (GALs). Thus, these studies indicate that keratinocytes generate the lysosomes in ATF6-dependent manner from Golgi during calcium-induced differentiation process.

**Figure 7.**
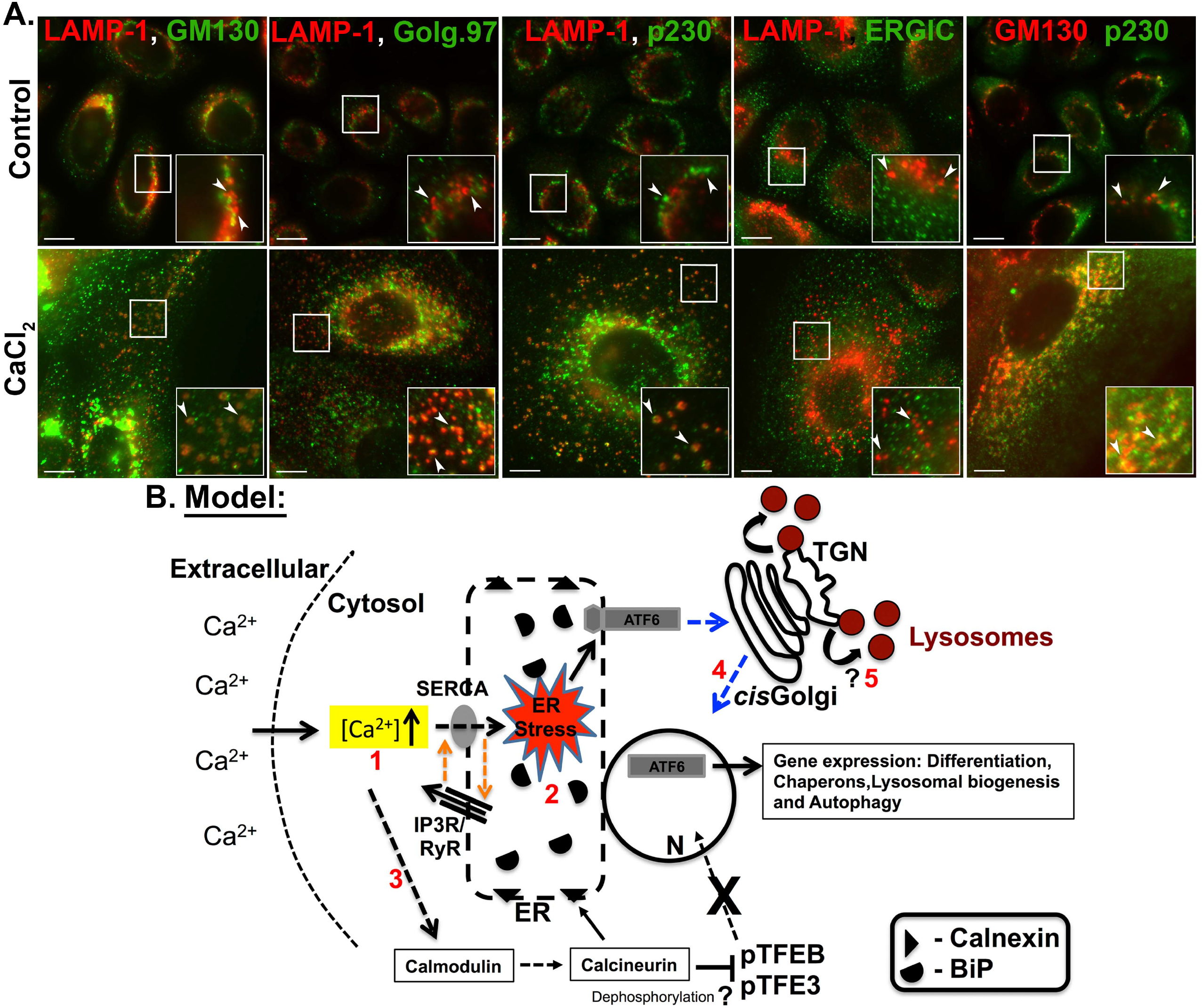
keratinocyte differentiation induced lysosomes are associated with Golgi tethering proteins and the model illustrating the mechanism of keratinocyte differentiation and lysosome biogenesis. (A) IFM analysis of control and differentiated keratinocytes for the localization of Golgi-associated protiens with respect to the lysosomes. Arrowheads are point to the colocalization of Golgi proteins with lysosomes. The insets are a magnified view of the white boxed areas. Scale bars, 10 μm. (B) Prolonged exposure of keratinocytes with CaCl_2_ increases intracellular calcium [Ca^2+^] levels (1) within 2 hr and possibly elevates ER stress (2) due to altered ER calcium refilling cycle/gradient (orange arrows). Cytosolic calcium levels likely increases the activity of calcium signalling moleucles calmodulin and calcineurin (3), but result in retention of MiT/TFE TFs to the cytosol. Enhanced ER stress promote the Golgi trafficking and processing of ATF6α in to an active UPR TF (4), which possibly initiates the cell differentiation and lysosome biogenesis including BiP levels. During this process, Golgi compartments merged and generate the enlarged globular lysosomes (5), which probably senses intracellular signaling and balances calcium concentration by acting as reservoir. ? indicate the pathway requires future investigation.

## Discussion

Cellular differentiation is one of the adaptive survival machanisms in response to extracellular stimuli. Epidermal keratinocytes of basal layer encounter both nutritional stress and ionic (calcium) stimuli that lead to a change in cell fate from proliferation to differentiation^2,64^ in addition to an increase in intracellular digestion through autophagy^8,9^. Several studies have reported that extracellular high calcium plays a key role in inducing keratinocyte differentiation^2^, but its role in upregulation of cellular clearance activity is poorly understood. Our *in vitro* model of keratinocyte differentiation using CaCl_2_ illustrated a dramatic increase in the number and size of functional Golgi-derived lysosomes (GALs), which possibly maintains the cellular digestion/macroautophagy. Moreover, our study demonstrated that an initial rise (early as 2 h) in cytosolic calcium possibly triggers ATF6 arm (but not IRE1 or PERK) of UPR that enhances lysosome biogenesis independently of mTOR and MiT/TFE TFs. Consistently, inhibition of ER/Golgi function, but not the lysosome/-associated signaling activity abolished the biogenesis of GALs and the differentiation of keratinocytes. Thus, our study provides the molecular mechanism of cellular homeostais during keratinocyte differentiation in response to extracellular high calcium.

How extracellular high calcium lead to GAL biogenesis? We predict that keratinocytes balance the calcium stress by maintaining the active calcium gradient between extracellular environment - cytosol - intraorganelles such as ER, lysosomes etc.^65,66^. Elevated calcium levels in the cytosol induces multiple calcium-dependent singalling pathways that may lead to the activation of apoptosis^67,68^. To escape the programmed cell death, cells are very likely to accommodate excess intracellular calcium in membrane bound organelles such as GALs in addition to ER and mitochondira, which protect the cell against the activation of multiple calcium-dependent signalling pathways. Several observations in our study support this hypothesis: (1) the cytosolic calcium level was lower in differentiated compared to proliferating keratinocytes; (2) inhibition of SERCA pump using Tg completely reduced the lysosome biogenesis and differentiation; (3) small molecule inhibition of calmodulin or calcineurin showed moderate effect on differentiation and (4) activation of ATF6 branch of UPR enhances the organelle biogenesis such as lysosomes and autophagosomes. In line with these results, a co-ordinated role between cell surface calcium pumps with ER SERCA in maintaining the low cytosolic calcium concentration has been implicated previously^69^. Thus, GAL biogenesis during keratinocyte differentiation is one of the possible mechanisms to nullify the nutritional and ionic stresses in the epidermis.

Studies have shown that differentiated keratinocytes in the upper layers of epidermis gradually lose intracellular organelles by increasing macroautophagy^8,9^. In turn, the enhanced autophagy requires functional lysosomes to maintain the turn over of intracellular digestion. But it is unclear how these lysosomes are produced during keratinocyte differentiation. Our study provide a model in which initial intracellular calcium rise (within 2 h) creates ER stress (step 1 and 2), which then activates the ATF6 arm of UPR and generates active ATF6 TF (step 4). ATF6 localize to the nucleus and possibly upregulate the genes for differentiation, lysosome biogenesis, autophagy etc. In parallel, cells promote the generation and dispersal of functional lysosomes from Golgi, called GALs (step 5) to reduce the extracellular calcium induced ER stress. Interestingly, cytosolic calcium known to enhance the activity of calmodulin^57^ and calcineurin and the later molecule known to dephosphorylate the MiT/TFE TFs^30,35^. We have hypothesized that the calcineurin activity is very likely to be diverted towards calnexin (an ER chaperon)^58^ (requires future investigation), by which the phosphatase was unabled to activate the TFEB/TFE3 duing keratinocyte differentiation (step 3). Alternatively, active mTOR sequesters TFEB/TFE3 TFs on lysosome membranes during differentiation and high mTOR activity was probably nullified the calcineurin activity. Consistently these TFs unable to localize to the nucleus and retained in the cytosol of differentiated keratinocytes. Thus the process of lysosome biogenesis follow a non-canonical secretory pathway involving ER stress rather than mTOR-MiT/TFE TFs-dependent classical biogenesis process^27–29^. Several results of our study support this pathway: (a) increased ATF6α TF levels and its localization to the nucleus over time; (b) retention of MiT/TFE TFs to the cytosol due to enhanced mTOR activity; (c) localization of pmTOR to the lysosomes; (d) association of Golgi tethering factors to lysosomes; and (e) GALs possess all the characteristics of endocytic pathway derived lysosomes such as acidity, proteolytic acitivity, localization of lysosome-specific GTPases and membrane proteins. Additionally, our model of lysosome biogenesis support the balance required for the turn over of enhanced autophagosomes/autophagy flux observed during keratinocyte differentiation.

Does lysosome biogenesis is essential for keratinocyte differentiation or it is a cause and effect of the differentiation program? Several studies have reported that autophagy is the key process known to effect the epideraml differentiation and skin architecure/function^8,9,31,70^. The molecular link between these processes is critically unknown. Recent studies have shown that mTOR dependent MiT/TFE TFs such as TFEB and TFE3 regulate both lysosome biogenesis and autophagy by binding to the CLEAR element network^50^. Initially, we predicted that MiT/TFE TFs including MITF possibly regulate keratinocyte differentiation and lysosome biogenesis/autophgy. To our surprize, none of the MiT/TFE family members enhanced their localization to the nucleus despite their significantly upregulated protein levels during calcium-induced differentiation of keratinocytes. This is possibly due to the upregulated activity and localization of mTOR to the lysosomes, which phosphorylates MiT/TFE TFs and causes the cytosolic retention of these factors during keratinocyte differentiation^27–29^. Alternatively, activated AKT signalling observed in differentiated cells may also phosphorylate TFEB^71^. In contrast, autophagosome number and their turnover significantly increased in the differentiated cells. These results led us to identifying a non-canonical pathway involving ER stress that regulates the lysosome biogenesis and keratinocyte differentiation. Chemical inhibition of pathways revealed that modulation of ER stress or Golgi function by using Tg/4-PBA or brefeldin A respectively, completely abolished the keratinocyte differentiation and lysosome biogenesis (Figure 5 and Table 1), however, alteration of lysosome acidity using bafilomycin A1 had no effect on these processes (only D_L_ is reduced, see Figure 5 and Table 1). Thus, our studies provide evidence that ER stress is essential for keratinocyte differentiation, which regulates the biogenesis of GALs to compensate the increased autophagic flux. Additionally, these GALs may also regulate the epidermal barrier function that requires ceramide secretion from these organelles (investigated in future). Overall, ionic calcium in the basal layer of epidermis initiate the differentiation as an apaptive mechanism to compensate the nutrient stress observed in the skin layers. Additonally, these studies will help in understanding the epidermal homeostasis or architecture, which is defective in several skin diseases including psoriasis and atopic dermatitis.

## Materials and methods

### Cell culture, differentiation, transfection and durg treatment

Neonatal human epidermal keratinocytes (NHEK) were purchased from Lonza or Invitrogen. Early passaged cells (less than 7) were maintained in EpiLife serum free medium (contains 60 μM CaCl_2_) supplemented with human keratinocyte growth supplement (HKGS). Cells were plated on 0.002% collagen coated surface for better adherence and then incubated at 37 °C with 10% CO_2_. For differentiation, keratinocytes at a confluency of 50 – 60 % were supplemented with 2 mM CaCl_2_ for 2 – 9 days and changed the medium every 24 h. We observed more than 70% (see Table 1) cells showed differentiation characteristics at 48 h and were used for all experiments. DNA vectors were transfected into the cells by using Lipofectamine 2000 (Invitrogen) according to the manufacturer’s protocol. Cells were fixed, immunostained and imaged as described previously^72^. For drug treatment, cells were incubated with respective drug concentrations (listed in Table 1, obtained from the literature) along with 2 mM CaCl_2_ for 48 h, fixed and then stained. Note that cells were replenished with their respective medium at 24 h. Moreover, we used lower drug concentrations to avoid reversible effect of the compound within 48 h. To equivalize the time, cells were treated with drugs throughout the 48 h time period. Half-cell length (CL_H_) as parameter to measure differentiation and dispersion of lysosomes (D_L_) as parameter to measure the lysosome biogenesis were quantified (see below) in μm (from the nucleus to cell periphery) and then compared with CaCl_2_-treated cells. For lysotracker staining, cells on glass coverslips were incubated with 50 – 75 nM of lysotracker in growth medium for 30 min at 37 °C in a incubator maintained at 10% CO_2_. Similarly for DQ-BSA internalization, cells were incubated with 10 μg/ml of DQ-BSA in a complete medium for 2 h, washed once with 1X PBS and then incubated in plain medium for 4 h. Finally, cells were fixed with 3% paraformaldehyde solution, stained and imaged.

### Epidermal skin grafts and immunohistochemistry

Epidermal foreskin grafts or human skin grafts were obtained as waste discard post cosmetic surgery with informed patient consent and local ethical committee approval. These grafts were embeded in paraffin blocks, sectioned into 5 μm thick slices using microtome (Leica RM 2155) and then placed on poly-L-lysine coated slides (Sigma-Aldrich). Sections were deparaffinized using xylene followed by rehydration sequencially with ethanol (100%, 90%, 70%) and then with distilled water. Slices were further treated with 10 mM sodium citrate buffer pH 6.1 and then incubated in 1XPBS for 45 min. followed by an incubation in 1.5% blocking serum in PBS (Santa Cruz Biotechnology, sc-2043) for 1 h at room temperature under moist condition. After the removal of excess blocking serum, the sections were immunostained with indicated antibodies (1:20 dilution in blocking serum) for overnight at 4 °C. Slides were rinsed with 1XPBS containing 0.05% Tween 20 for 3 times (5 min. each) and then stained with respective secondary antibodies for 1 h at room temperature. Finally, slides were washed and then imaged.

### Immunofluorescence microscopy and image analysis

Cells on coverslips were fixed with 4% formaldehyde (in PBS) and then stained with primary antibodies followed by the respective secondary antibodies as described previously^72^. In some experiments, cells on coverslips were internalized with lysotracker or DQ-BSA, fixed, immunostained and imaged. Bright-field (BF) and immunofluorescence (IF) microscopy of cells was performed on an Olympus IX81 motorized inverted fluorescence microscope equipped with a CoolSNAP HQ2 (Photometrics) CCD camera using 60X (oil) U Plan super apochromat objective. Acquired images were deconvolved and analyzed using cellSens Dimension software (Olympus). The colocalization between two colors was measured by selecting the entire cell excluding the perinuclear area and then estimated the Pearson’s correlation coefficient (r) value using cellSens Dimension software. The average *r* value from 10-20 cells was calculated and then represented as mean±s.e.m. Note that the maximum intensity projection of undeconvolved Z-stack images were used for the measurement of *r* values. Analyzed images were assembled using Adobe Photoshop. Half-cell length (in μm, labeled as CL_H_) was measured as the maximum distance between nucleus and the cell periphery using cellSens Dimension software. Likewise, length and width of the nucleus was measured (in μm) by placing the scale bar along the diameter of the nucleus. In Figure 5, half-cell length (in μm) was measured by masking the BF images with LAMP-1 staining (except bafilomycin A1 condition) to distinguish the cell border. In parallel, the distribution of lysosomes (in μm, labeled as D_L_) in each condition was measured independently from the nucleus to cell periphery (the longest possible distance) using LAMP-1 stained IFM images. Note that the pararameter CL_H_ was measured independently in bafilomycin A1 treated cells. Averages of CL_H_ and D_L_ (in μm) in each condition were calculated from approximately 60 – 80 cells (*n*=2-3), indicated in the Figure 5 and Table 1 (mean ± s.e.m.), and then plotted using GraphPad Prism software. Using the half-cell length parameter, the percentage of undifferentiated cells (approximately to the maximum size of control cells, < 19 μm) in a given drug treatment/condition was calculated from two or three different experiments and then indicated in Table 1. In Figure 1E, the percentage of cells showing perinuclear or peripheral distribution of LAMP-1-positive compartments was visually quantified (representing the pattern similar to Figure 1A, 4^th^ panel) from three different experiments and then plotted. Similarly, number of GFP-LC3 puncta were visually quantified from three different experiments and then plotted.

### Statistical analysis

All statistical analyses were done using GraphPad Prism 5.02 and the significance was estimated by unpaired Student’s *t* test. *, *p*≤0.05; **, *p*≤0.01; ***, *p*≤0.001 and ns, not significant.

## Abbreviations

4-PBA: 4-Phenylbutyric acid
AMPK - AMP: -activated protein kinase
AKT/PTB: protein kinase B
ATF6: activating transcription factor 6
BF: bright-field
Calcium: CaCl_2_
ER: endoplasmic reticulum
IFM: immunofluorescence microscopy
PI3K: phosphoinositide 3-kinase
TFEB: transcription factor EB
Tg: thapsigargin
mTORC1: mammalian target of rapamycin complex 1
UPR: unfolded protein response

## Acknowledgements

We thank A. Gupta, S. Nag and U. Krishnan for their technical help. This work was supported by a Wellcome Trust-DBT India Alliance Senior Fellowship (500122/Z/09/Z to S.R.G.S.); Unilever Industries Pvt. Ltd., Bangalore; CEFIPRA Project (4903-1 to S.R.G.S. and Graca Raposo); DBT-RNAi task-force (BT/PR4982/AGR/36/718/2012 to S.R.G.S.); IISc-DBT partnership program (to S.R.G.S.) and S.M. was supported by DBT-IISc partnership postdoctoral fellowship.

## Author contributions

S.M. designed and performed all the experiments in this study and participated in manuscript writing. S.S.D. performed standardization of keratinocyte differentiation experiments and transcriptional analysis of lysosomal genes. R.S. performed immunostaining of epidermal skin grafts. S.P. carriedout lysosomal enzyme activity assay. M.M.S. maintained primary keratinocytes. A.M. provided several reagents and scientific support throughout the project period. S.R.G.S. oversaw the entire project, coordinated and discussed the work with coauthors, and wrote the manuscript.

## Materials and Correspondence

For material request contact

## Online supplemental information

Details of materials, reagents and few experimental procedures are described in the Supplemental information. Supplementary Figure 1 (related to Figures 1 – 3) shows CaCl_2_ but not CaCO_3_ induces keratinocyte differentiation that alters nuclear size; however, the transcription of genes or characteristics of endocytic organelles remain unchanged. Supplementary Figure 2 (related to Figures 6 – 7) shows expression of UPR regulated factors and the localization of Golgi-tethering factors to lysosomes. Supplementary Table 1 lists the oligo sequences used for transcript analysis.

Primary keratinocytes were treated with the indicated compounds (target and their concentrations were listed) along with CaCl_2_ for 48 h, fixed, stained with LAMP-1 and then imaged (see Figure 5). In condition 11, cells were treated with CaCl_2_ for 24 h and then incubated with 4-PBA. In condition 12, cells were pre-incubated with 4-PBA for 24 h and then replaced with CaCl_2_. Keratinocyte differentiation and lysosome biogenesis were quantified as half-cell length (CL_H_) and lysosome dispersion (D_L_), respectively in the indicated treatments (see Figure 5) as described in the materials and methods. Both CL_H_ and D_L_ were measured as μm from the nucleus towards cell surface (approximately 60-80 cells, *n*=2-3). Average CL_H_ and D_L_ (in μm) for each treatment were indicated (mean ± s.e.m.). The percentage of undifferentiated cells (having CL_H_ < 19 μm, maximum length observed in proliferative keratinocytes) was calculated from the data presented in Figure 5B and 5D and listed separately in the table.

## Supplementary Information

### Materials and methods

#### Reagents and antibodies

All chemicals and reagents were purchased either from Sigma-Aldrich (Merck) or ThermoFisher Scientific (Invitrogen). Torin 1 from Tocris Bioscience was purchased. Tissue culture reagents such as EpiLife medium, HKGS (Human Keratinocyte Growth Supplement), Trypsin neutralizer solution and reagents including LysoTracker Red DND-99, DQ-BSA, Fluo-4 NW were obtained from ThermoFisher Scientific (Invitrogen).

The following commercial polyclonal and monoclonal antisera were used (m, mouse; h, human and r, rat proteins). Anti-mLIMPII (ab16522) was from Abcam; anti-rGM130 (610822) and anti-hp230 (611281) were from BD Biosciences; anti-pAKT (9271), anti-Beclin-1 (3495); anti-BiP (3177), anti-Calnexin (2679), anti-CHOP (2895), anti-4E-BP1 (9452), anti-EEA1 (3288), anti-IRE1α (3294), anti-PERK (5683), anti-Rab9 (5118), anti-Raptor (2280); anti-Rictor (2114); anti-LC3A/B (4108), anti-pS6K (T389; 9234), anti-SQSTM1 (p62; 5114), anti-TFEB (4240), anti-mTOR (2983) and anti-p-mTOR (S2448; 5536) were from Cell Signalling Technology; anti-hLAMP-1 (H4A3) and anti-hLAMP-2 (H4B4) were from Developmental Studies Hybridoma Bank; anti-hATF-6α (sc-22799), anti-hInvolucrin (sc-21748), anti-Keratin14 (sc-53235), anti-hMITF (sc-10999) and anti-mXBP-1 (sc-7160) were from Santa Cruz Biotechnology; and anti-β-actin (A5441), anti-hGBA (G4171), anti-HA (H3663), anti-Histone H3 (H9289), anti-TFE3 (HPA023881) and γ-tubulin (GTU88; T6557) from Sigma-Aldrich. Antisera to Golgin97, ERGIC (Michael S. Marks, University of Pennsylvania, Philadelphia, USA) and STX13 (Andrew Peden, University of Sheffield, Sheffield, UK) were obtained as gift from respective laboratories. All secondary antibodies were either from Invitrogen or Jackson Immunoresearch.

#### Plasmids

Arl8b-GFP – Human Arl8b was PCR amplified from cDNA derived from HeLa cells, digested and subcloned into XhoI and BamHI sites of pEGFP-N1 vector (Clontech). TFEB-GFP – Human TFEB was PCR amplified from HeLa cells cDNA, digested and subcloned into SalI and BamHI sites of pEGFP-N3 vector (Clontech). HA-Rab7A was a kind gift from Michael S. Marks, University of Pennsylvania, Philadelphia, USA and GFP-rLC3 (21073) was obtained from Addgene.

#### Lysosomal enzyme (Glucerebrosidase) activity assay

Keratinocytes were seeded at 70 – 80% confluence in black 96 well flat bottom black plate (Corning) and performed the intact cell lysosomal β-glucosidase assay as described previously^S1^. Briefly, post treatment, cells were washed twice with 1XPBS and then incubated with 50 μl of 3 mM MUD (4-methyl umbelliferyl-β-D-glucopyranoside, made in 0.2 M sodium acetate buffer pH 4.0) at 37 °C for 3 h. Further, the assay was stopped by adding 150 μl of 0.2 M glycine buffer pH 10.8 and then measured the fluorescence intensity of liberated 4-methylumbelliferone (excitation at 365 nm and emission at 445 nm) using Tecan multi-mode plate reader (Infinite F200 Pro). In parallel, cell density in the duplicated wells was calculated using Resazurin (Sigma-Aldrich)-based cell viability assay. Here, 10 μl of Resazurin (10 mg/ml made in 1XPBS) in 90 μl of medium was added to the cells and then incubated at 37 °C for 6 – 7 h. The fluorescence intensity was measured at 530 nm excitation and 590 nm emission using multi-mode plate reader (Tecan). Finally, the lysosomal enzyme activity was normalized with their respective cell viability values and then plotted.

#### Measurement of intracellular calcium levels

Primary keratinocytes were seeded at 70 – 80% confluence in black 96 well flat bottom plate (Corning) and incubated with CaCl_2_ for different time intervels or treated with Tg or Torin 1 or EGTA (control) in combination with CaCl_2_ for 6 h or 48 h. Intracellular free calcium was measured using Fluo-4 NW calcium assay kit (Molecular probes, F36206). Briefly, Fluo-4 NW dye was added to the cells 2 – 6 h prior to the end of time point. The total cellular fluorescence was measured at 516 nm with an excitation of 494 nm using Tecan multi-mode plate reader (Infinite F200 Pro). The emission fluorescence intensity values were normalized with the cell numbers measured by Resazurin (Sigma-Aldrich)-based cell viability assay. The fold change in fluorescence intensity between the treatment and control was measured and then plotted.

#### Cell surface expression using flow cytometry

Cell surface expression of LAMP-1 and LAMP-2 was measured as described previously^S2^. Briefly, cells were harvested, washed once with 1XPBS, suspended in ice-cold growth medium (supplemented with 25 mM HEPES pH 7.4) containing anti-LAMP-1, anti-LAMP-2 or anti-Tac (7G7.B6, ATCC, as negative control) and incubated on ice for 30-45 min. Cells were washed, suspended in medium containing respective Alexa 488-conjugated secondary antibodes and then incubated on ice for 30-45 min. Finally, cells were washed, suspended in ice-cold FACS buffer (5% FBS, 1 mM EDTA and 0.02% sodium azide in PBS) and measured the fluorescence intensity using FACS Canto (BD biosciences). Data was analyzed using FlowJo (Tree Star) software and then plotted the mean fluorescence intensity (MFI) as described previously^S2^.

#### Transcript analysis by semi-quantitative PCR

Keratinocytes grown in a 35 mm dish was subjected to RNA isolation using GeneJET RNA Purification kit (ThermoScientific). The cDNA was prepared from total RNA by using a cDNA synthesis kit (Fermantas). Gene transcripts were amplified (Bio-Rad S1000 Thermal Cycler) using an equal amount of cDNA from each condition and the gene specific primers (listed in Supplementary Table 1). In all experiments, GAPDH or 18S rRNA was used as loading control. Band intensities were measured, normalized with the loading control, quantified fold change with respect to the control (three independent experiments) and then indicated in the figure.

#### Nuclear-cytosolic fractionation

Separate dishes of cells were used for cytosolic and nuclear extract preparation. Before use, cells were washed twice with 1XPBS. For cytosolic extraction, cells in a dish was added with 400 μl buffer (10 mM HEPES-KOH pH 7.5, 3 mM MgCl¾ 40 mM KCl, 1.0 mM DTT, 0.1 mM PMSF, 0.3% NP-40 and protease inhibitor cocktail) and incubated on ice for 10 min. Cells were scrapped, collected, incubated on ice for additional 15 min, centrifuged at 15000 rpm for 30 min and separated the supernatant. Similarly, for nuclear extraction, 500 μl of buffer (10 mM HEPES-KOH pH 7.5, 10 mM KCl, 1.0 mM DTT, 0.1 mM PMSF and protease inhibitor cocktail) was added to the dish of cells and incubated on ice for 10 minutes. Further, 0.3% NP-40 was added to the dish and then incubated for 10 min on ice. Cells were scrapped, collected, centrifuged at 15000 rpm for 5 min and discarded the supernatant. Pellet was suspended in 200 μl of buffer (20 mM HEPES-KOH pH 7.5, 400 nM NaCl, 1.0 mM DTT, 0.1 mM PMSF and protease inhibitor cocktail), incubated for 10 min on ice, centrifuged at 15000 rpm for 30 min and separated the nuclear lysate.

#### Immunoblotting

Cell lysates were prepared in RIPA buffer and then subjected to immunoblotting analysis as described previously^S2^. Immunoblots were developed with Clarity Western ECL substrate (Bio-Rad) and the luminescence was captured using Image Lab 4.1 software in a Bio-Rad Molecular Imager ChemiDoc XRS+ imaging system, equipped with Supercooled (-30°C) CCD camera (Bio-Rad). Protein band intensities were measured, normalized with loading control, quantified the fold change with respect to control and then indicated in the figure.

### Supplementary figures

**Supplementary Figure 1.**
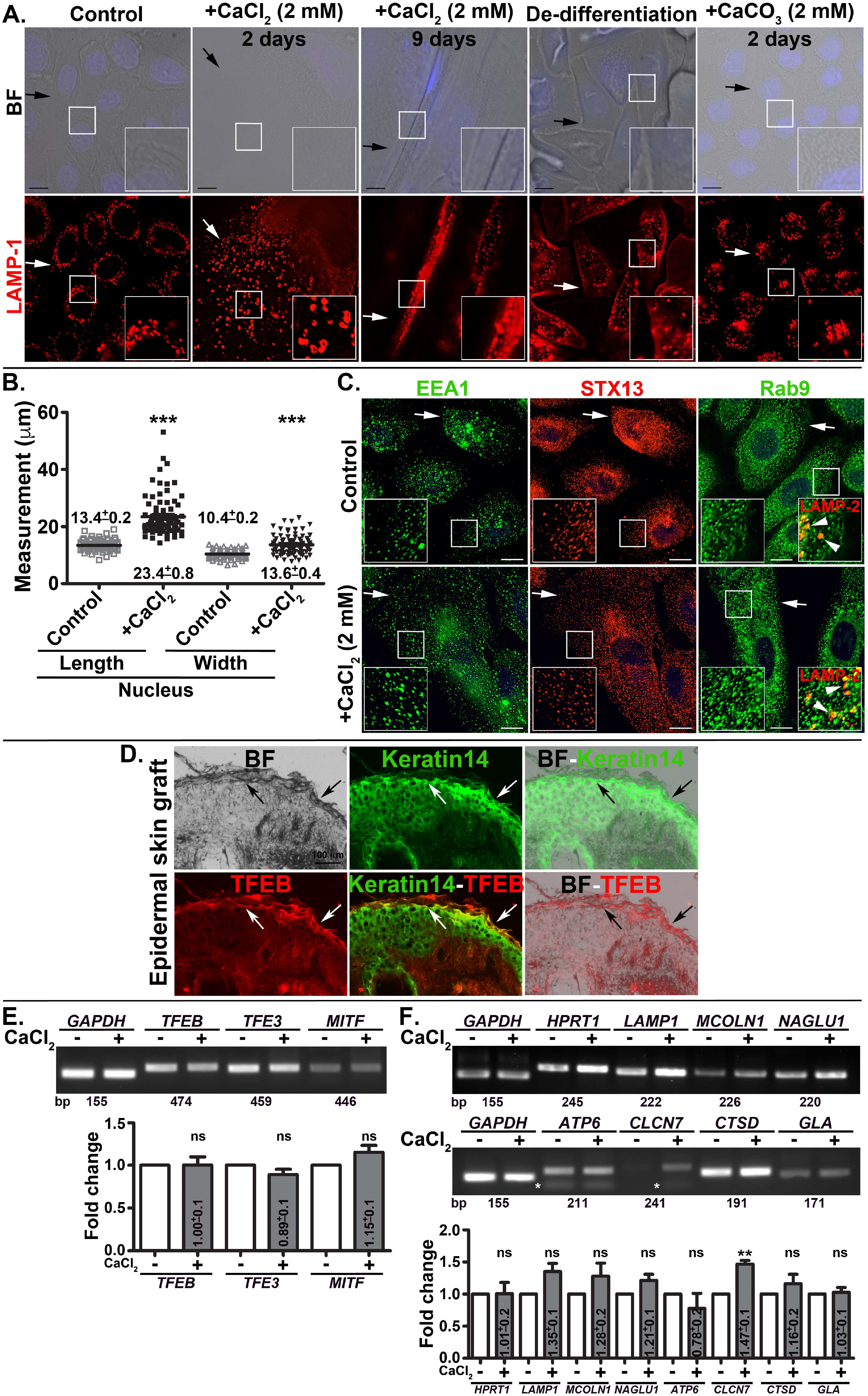
Keratinocyte differentiation is induced with CaCl_2_, but not with CaCO_3_ and the process is not reversible. Differentiation of cells alters the nuclear size, but not the endocytic organelles and transcription of genes. (A, C) BF and IFM image analysis of control, CaCl_2_ or CaCO_3_-incubated cells for 2 or 9 days. In panel 3, cell showed the characteristics of terminal differtiation. In panel 4, CaCl_2_ was removed and then grow the cells for two days (de-differentiation) in regular medium. Cells were immunostained for endocytic and lysosomal proteins as indicated. Black arrows point to the cell size and white arrows indicate the distribution of early/recycling endosomes and lysosomes. Arrowheads point to LAMP-2-positive lysosomes with respective to Rab9. The insets are a magnified view of the white boxed areas. Nuclei are stained with Hoechst33258. Scale bars, 10 μm. (B) Measurement of nuclear length and width (in μm) in keratinocytes. Approximately 100 nuclei (*n*=3) from each condition were measured and then plotted. Average length/width (mean ± s.e.m.) of the cell was indicated in the graph. ***, *p*≤0.001. (D) BF and IFM of epidermal skin graft that was immunostained for keratinocyte differentiation maker keratin14 and TF TFEB. Black arrows point to the cornified outermost layer located above the immunostained keratinocyte layer. White arrows indicate the differentiated keratin14-positive keratinocytes that showed enhanced TFEB expression. Scale bars, 100 μm. (E, F) Semiquantiative PCR analysis of control and differentiated cells to measure the expression of MiT/TFE TFs and various lysosomal biogenesis genes. Expected size (in base pairs) of gene was indicated. Band intensities were quantified, normalized with loading control (GAPDH) and plotted the average fold-changes (mean ± s.e.m., *n*=3). * indicates non-specific PCR bands. **, *p*≤0.01 and ns, not significant.

**Supplementary Figure 2.**
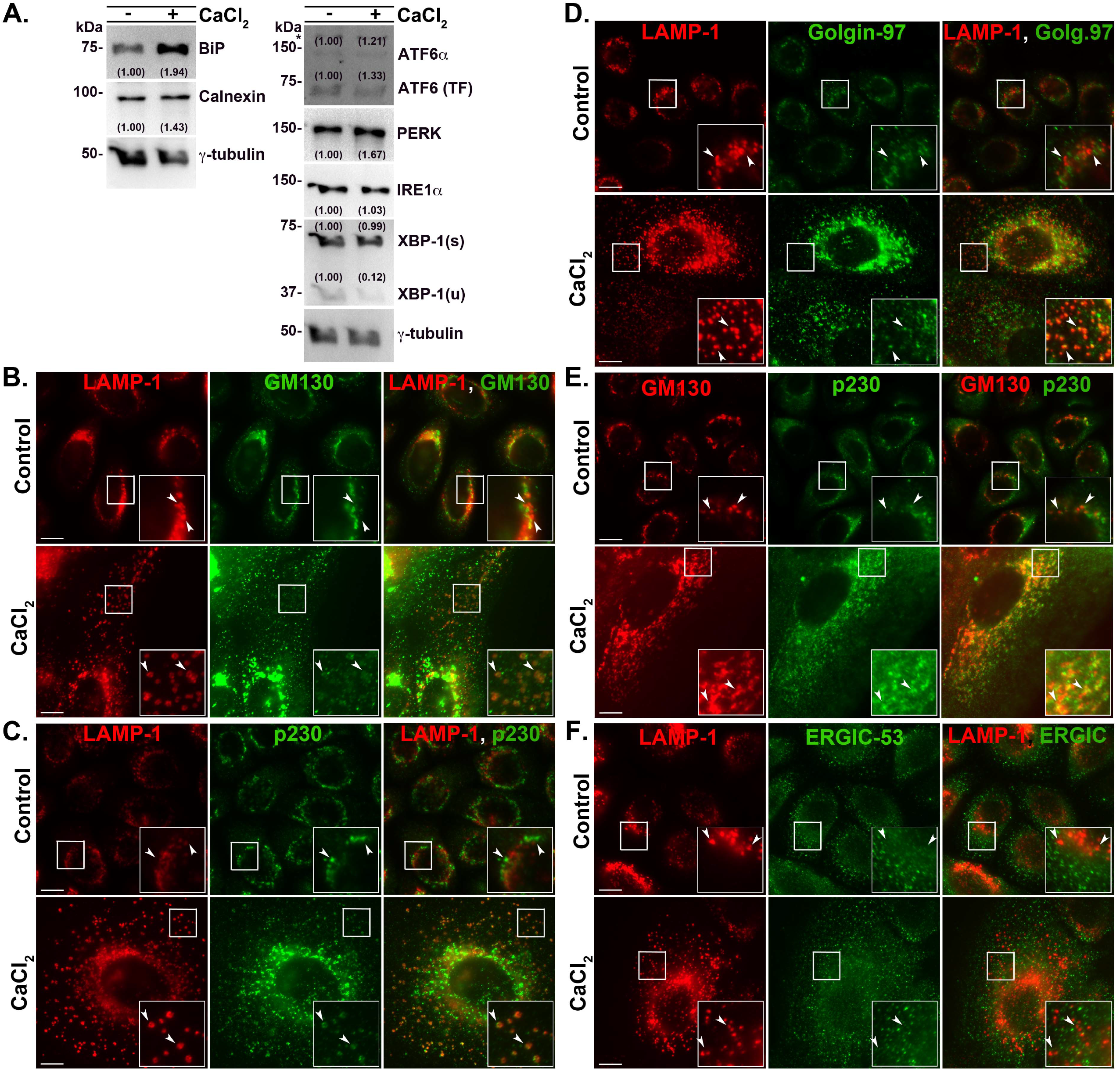
(A) Immunoblotting analysis of the UPR regulated proteins and their downstream factors. Band intensities were quantified, normalized with loading control (γ-tubulin) and mentioned the fold-change on the gels. (B) IFM analysis of control and differentiated keratinocytes for the localization of Golgi-associated proteins with respect to the lysosomes. Arrowheads are point to the colocalization between lysosomal protein with Golgi-associated or ER-Golgi proteins. The insets are a magnified view of the white boxed areas. Scale bars, 10 μm.

**Supplementary Table 1.**
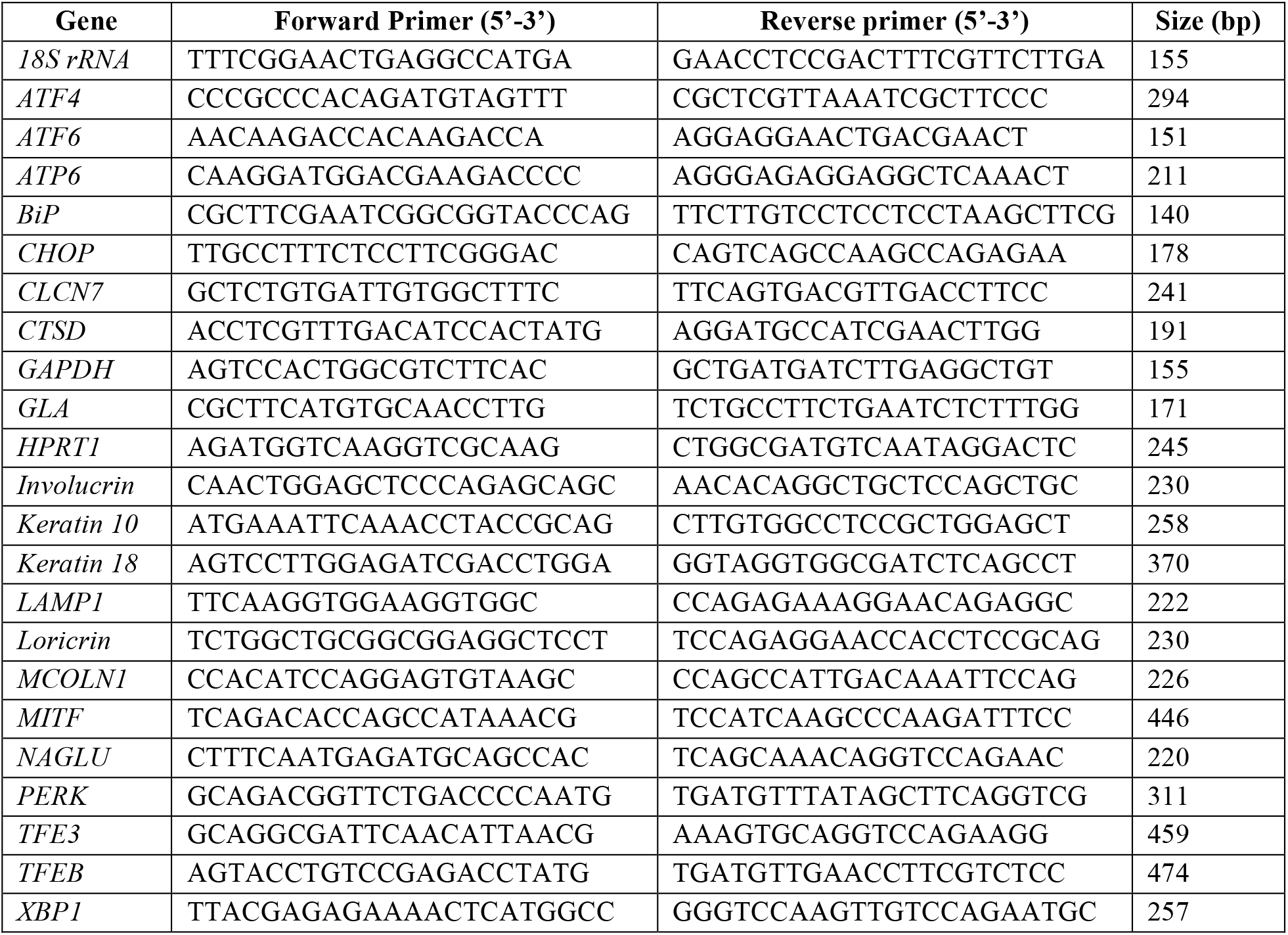
List of primers used for transcript analysis

